# MICROBIOTA AT THE CROSS ROAD OF DIET AND HEALTH: HOW DIETARY FATS SHAPE BACTERIAL LANDSCAPE AND INFLUENCE GLOBAL HEALTH

**DOI:** 10.1101/2024.07.08.602483

**Authors:** Néstor D. Portela, Natalia Eberhardt, Gastón Bergero, Yanina L. Mazzocco, Maria P. Aoki, Cristian A. Galván, Roxana C. Cano, Susana A. Pesoa

## Abstract

Host-gut microbiota (GM) interactions play a pivotal role in shaping the delicate balance between health and disease within the human body. The impact of dietary factors, specifically high fat content diets on GM composition has been widely demonstrated. We have previously shown that the constant and sustained administration of Omega-3 fatty acids induced specific changes in GM composition, modulating the immune metabolic response of visceral adipose tissue (VAT) in our mouse model of obesity. We now set out to determine if this effect is Omega-3 dose-dependent. To achieve this, C57BL/6J(B6) mice were fed for 24 weeks with three diets, two with medium content total fat, but different Omega-3 content and a control diet.

GM composition, metabolic biomarkers and immune cells in VAT were analyzed. A distinctive segregation of GM composition, a significantly higher proportion of regulatory T cells (CD45+CD4+FoxP3+), Omega-3 dose dependent and increased levels of leptin and cholesterol with no differences in adiponectin values were found in fat fed groups. Simple mediation analyses revealed significant associations between the microbial profile and immunometabolic regulation. To remark is the capacity of *Lachnospiraceae UCG- 001* to modulate levels of leptin, glucose, and cholesterol through the stimulation of CD45+CD4+FOXP3+IL10+ cells. Our findings suggest a modulatory effect of omega-3 fatty acids on the microbiota, the metabolism, and the immunoregulatory capacity of VAT, supporting the hypothesis that alteration of the GM composition by omega-3 fatty acids may be a promising approach in managing obesity and associated metabolic diseases.

## 1. Introduction

Obesity is a complex disease associated with a range of adverse health consequences, including cardiovascular diseases, insulin resistance, Type 2 Diabetes Mellitus (T2DM), hypertension, and certain types of cancer (**1**). This condition is the result of a sustained positive energy balance, leading to excessive accumulation of VAT which becomes dysfunctional as the pathology progresses, driving to chronic inflammation and altering its role in the regulation of overall metabolism (**2,3**). Environmental factors, such as diet and sedentary lifestyle, play a crucial role in the development of obesity. Additionally, numerous articles have been published linking specific changes in the GM to the development of obesity and its associated disorders (**4–6**). The GM is involved in many functions, including nutrient absorption, protection of intestinal mucosal integrity, regulation of immune responses, being a central regulator of host metabolism (**7,8**). Furthermore, multiple articles have underscored the importance of diet in regulating the composition of the GM (**9–11**).

It is well established that both the GM structure and function are dynamic and strongly affected by diet nutrients, such as the content and composition of lipids (**12–15**). As a consequence, dietary lipids influence host physiology through interactions with the GM, although the mechanisms involved are not fully defined (**16**).

On the other hand, the infiltration of pro-inflammatory macrophages and CD8+ T cells into adipose tissue is a crucial component that amplifies persistent inflammation (**17,18**). Simultaneously, the decrease in anti-inflammatory immune cells, such as M2 macrophages, eosinophils, and regulatory T cells (Tregs), exacerbates the immunological imbalance (**19**). The dysregulation of adipocytokines, crucial molecules in metabolism modulation, along with endoplasmic reticulum stress, also contributes significantly to the inflammatory cascade (**20,21**). In this context, diverse experimental models of obesity have highlighted the importance of M2 macrophages due to their specifically anti-inflammatory role in this process. Likewise, Tregs, recognized for their function in immune tolerance, show a marked reduction in epididymal fat of obese animals compared to lean controls, and this decrease is closely related to insulin resistance (**22**). On the other hand, it has been reported that adipose tissue macrophages (ATMs) may promote the differentiation and proliferation of Tregs in VAT in vivo (**23**).

We have previously demonstrated that the constant and sustained administration of Omega-3 fatty acids induced specific changes in GM composition, modulating the immune metabolic response of adipose tissue in our mouse model of obesity. We now set out to determine if this effect is dose-dependent, making special focus on changes in the regulatory T cell abundance in VAT.

## 2. Materials and Methods

### 2.1. Animals and Experimental Design

Male C57BL/6J (B6) mice (6-weeks-old) were purchased from the Facultad de Ciencias Veterinarias of the Universidad Nacional de La Plata, Buenos Aires, Argentina. All animals were housed in isolation rooms at the Animal Facilities of the Facultad de Ciencias Químicas of the Universidad Católica de Córdoba.

The mice were randomly divided into three groups of four mice each and adapted to the experimental conditions for two weeks. The mice were then fed a low-fat diet, a “Control Diet” (D1): (4% fat and an Omega-3 fatty acid content of 0.8 g per 100 g of food), or Obesity-Inducing Diet composition - High content of Omega-3 (D2): (11% fat and an Omega-3 fatty acid content of 3.3 g per 100 g of food) or Obesity-Inducing Diet composition - Low content of Omega-3 (D3): with a medium fat content (11% fat and an Omega-3 fatty acid content of 1.6 g per 100 g of food) over a period of 24 weeks. All groups were maintained under a standard light cycle (12 h light/dark), with free access to water and food, and temperature conditions (21 ± 2 °C). The food was replaced every two days to keep it fresh.

This research has the authorization of the Institutional Committee for the Care and Use of Laboratory Animals— CICUAL-FCQ-Universidad Nacional de Córdoba, Córdoba, Argentina. Resolution N° 939, EXP-UNC: 0023836/2018. The environmental conditions in the facility were set according to CICUAL Guidelines. All animal procedures were performed in accordance with the guidelines of Directive 2010/63/EU of the European Parliament on the protection of animals used for scientific purposes.

### 2.2. Food Composition

Control Diet composition (D1): 4% fat, 26% protein, 52% carbohydrate, 8% crude fiber, and 10% total minerals (GEPSA Pet Foods, Pilar, Argentina).

Obesity-Inducing Diet composition - High content of Omega-3 (D2): 11% fat, 27% protein, 48% carbohydrate, 6% crude fiber, and 8% total minerals (Purina Nestlé).

Obesity-Inducing Diet composition - Low content of Omega-3 (D3): 11% fat, 28% protein, 4.5% crude fiber, and 7.5% total minerals (TIT Can Gross).

The lipid composition was as follows: D1, Total fat content: 4.0%, Omega-3 fatty acids: 20.3%, omega-6 fatty acids: 15.7%, Absolute Omega-3 fatty acids content: 3.3 g per 100 g of food, Absolute Omega-6 fatty acids content: 0.6 g per 100 g of food and Omega-3/omega-6 ratio: 1.3;

D2, Total fat content: 11.0%, Omega-3 fatty acids: 30.1%, omega-6 fatty acids: 8.7%, Absolute Omega-3 fatty acids content: 3.3 g per 100 g of food, Absolute Omega-6 fatty acids content: 1.0 g per 100 g of food and Omega-3/omega-6 ratio: 3.3.

D3, Total fat content: 11.0%, Omega-3 fatty acids: 20.3%, omega-6 fatty acids: 15.7%, Absolute Omega-3 fatty acids content: 1.6 g per 100 g of food, Absolute Omega-6 fatty acids content: 1.1 g per 100 g of food and Omega-3/omega-6 ratio: 1.4.

### 2.3. Food Intake Assay

To assess the acceptance of three diets, we carried out a consumption assay to measure food intake by mice. Briefly, we used individual mouse cages (n = 5/each diet) with one mouse each in the animal laboratory facility under the conditions cited above under Animals and Experimental Design. The initial amount of feed was weighed (5 g/cage to record daily feed intake). Consumption was measured by the differential weighing of the food at 0 and 24 h after feeding using an Ohaus digital scale model CS200, precision ± 0.1 g. Two independent experiments were performed to obtain an acceptable trend prior to the statistical analysis of the data (p-value < 0.05 was considered statistically significant) (**24**)

### 2.4. Evaluation of Obesity Development

The body weight of the mice was evaluated at 0, 4, 12, and 24 weeks post differential feeding start. Diet Induced Obesity (DIO) was defined by the cut-off point of the mean body weights of the mice at week 0 plus 3 standard deviations in all groups.

### 2.5. Analysis of Gut Microbiota Composition

The GM composition was studied in stool samples, collected individually in metabolic cages 24 weeks after the start of differential feeding. Feces were frozen immediately after collection and stored at −40°C until analysis.

#### 2.5.1. DNA Extraction

Stool samples were handled under a laminar flow hood using a sterile technique. Microbial DNA was isolated from 220 mg of stool using the QIAmp DNA Stool Mini Kit (Qiagen, Germantown, MD, USA) following the manufacturer’s standard protocol. The DNA was stored at −40 °C.

#### 2.5.2 16S rRNA Gene Sequencing and Taxonomic Identification

16S rRNA library preparation and sequencing was performed using Ion 16S Metagenomics Kit (Thermo Fisher Scientific, Carlsbad, CA, USA) on the Ion Torrent Personal Genome Machine (PGM) platform (Thermo Fisher Scientific, Carlsbad, CA, USA) according to the manufacturer’s instructions. A mock community dataset was generated from mixed bacterial genomic DNA from ATCC strains, including *Escherichia coli* ATCC 25922, *Staphylococcus aureus* ATCC 25923, *Pseudomonas aeruginosa* ATCC 27853, *Enterococcus faecalis* ATCC 29212, and *Streptococcus group B*; the latter was isolated and typified in the LACE Laboratory.

The sequence quality control, annotation, and taxonomical assignment were performed using the DADA2 v1.22.0 (**25**), phyloseq v1.38.0 (**26**), and microbiome v1.16.0 (**27**)packages in R software v4.1.2 (**28**) following the standard pipeline from demultiplexed fastq files. DADA2-formatted Silva Database Version 138.1–Updated 10 March 2021, was used for taxonomical assignment (**29**). Linear discriminant analysis Effect Size (LEfSe), was performed using microbiomeMarker package (**30**). Sequencing data are accessible in the National Center for Biotechnology Information (NCBI) database under BioProject accession number PRJNA1087685 (https://ncbi.nlm.nih.gov/bioproject/?term=PRJNA1087685) accessed on 20 March 2024.

### 2.6. Evaluation of Blood Metabolic Profile and Immune Cell Population in VAT

Twenty-four weeks after the start of differential feeding, all mice belonging to each group were anesthetized using inhaled FORANE (isoflurane). Blood samples were obtained after 8 h of fasting by cardiac puncture in heparinized tubes, centrifuged at 3000 rpm by 5 minutes, and the separated plasma was stored at −20 °C. The plasma glucose (Glu, mg/dL), triglyceride (TG, mg/dL), total cholesterol (TC, mg/dL), aspartate aminotransferase activity (AST, UI/L), alanine aminotransferase activity (ALT, UI/L), total proteins (g/dL) and Albumin (g/dL) levels were assessed in a ROCHE Cobas 8000 auto-analyzer (Roche Diagnostic). The adiponectin (ng/mL) and leptin levels (pg/mL) were quantified using an Adiponectin Mouse ELISA Kit (Abcam) and Leptin Mouse ELISA Kit (Invitrogen), respectively, following the manufacturer’s instructions. The mice were then sacrificed by cervical dislocation and the VAT was kept for studies of cell populations in stromal vascular fraction (SVF) as described below.

### 2.7. Isolation of the SVF from Adipose Tissue and Flow Cytometry

Mouse epididymal AT was processed by mechanical degradation and digested for 45 min at 37 °C with type 2 collagenase (0.8 mg/mL; Sigma) in Hanks’ Balanced Salt solution (pH = 7.4). After the addition of 3 vol. PBS containing 5% FBS and filtration of the digested tissue through nylon mesh (70 μm), the filtrate was centrifuged at 200× g. The SVF was recovered from the resulting supernatant (**31**). Red blood cells were separated by centrifugation at 500× g for 5 min, and the remaining cells were suspended in PBS and exposed to FcBLOCK (BD Biosciences) for 20 min. Five hundred thousand SVF cells were washed in ice-cold FACS buffer (PBS-2%FBS) and incubated with fluorochrome-labeled antibodies for 30 min at 4 °C. Different combinations of the following antibodies were used: PeCy5-labeled: anti-CD11b, PeCy7-labeled: anti-F4/80, APC-labeled: anti-CD45 and APCCy7-labeled: anti-CD4. Cells were permeabilized with BD Cytofix/Cytoperm and Perm/Wash (BD Biosciences) and a combination of the following intracellular antibodies were used: PerCp-Cy5.5 labeled: anti-FOXP3 and PE labeled: anti-IL-10. The cells were acquired on FACS Canto II (BD Bioscience). The results were expressed as the percentage of cells.

### 2.8. Statistical Data Analysis

The statistical analysis was conducted and visualized using R v4.1.2 software (**28**). The normality of variables was assessed using the Shapiro-Wilk test. Differential analysis between variables was conducted using the Student’s t-test or Wilcoxon test for pairwise comparisons between two variables, and ANOVA or Kruskal-Wallis test for the analysis involving more than two simultaneous variables. Pairwise comparisons using the Wilcoxon signed-rank test were performed if the Kruskal-Wallis test yielded a significant result, while the t-test was utilized in cases of significant results from the ANOVA test. The p-values were adjusted using the Bonferroni multiple-testing correction method. The correlation analysis between variables was executed using the corrplot package v0.92 (**32**).

For Lefse analysis, Linear Discriminant Analysis (LDA) scores of 3 and a p-value of <0.05 were considered significant. MicrobiomeMarket package v1.9.0 was used. All data were visualized using ggplot2 v3.4.0 (**33**) and ggpubr v0.5.0 (**34**). In all cases, a value of p < 0.05 was considered significant.

Alpha diversity metrics, including observed ASVs, Shannon, and Simpson indexes, as well as Beta diversity metrics (PCA and UniFrac, both weighted and unweighted), were computed based on the ASV table representing the relative abundances of bacterial taxa from the microbiome v1.6.0 R package (**27**). The relationship between diets and the overall microbiota composition was assessed using the Adonis test implemented through the Adonis function in the vegan v2.4.6 R package, as well as ANOSIM (**35**).

A mediation analysis was conducted using a linear regression model to explore the potential mediation effect of predictor variables on the predicted variables through mediating variables. The model was structured as follows:

Predictor Variables (X): Microbial genera obtained from Lefse analysis.

Mediating Variables (M): Immunological parameters evaluated in VAT.

Predicted Variables (Y): Main metabolic parameters evaluated.

The mediation model was defined with the following equations:

M=a×Predictor_Variable;

Y=c×Predictor_Variable+b×M;

Where:

a: represents the relationship between the predictor variable (Microbial genera) and the mediating variable (Immunological parameters).

b: represents the relationship between the mediating variable (Immunological parameters) and the predicted variable (Main metabolic parameters).

c: represents the direct relationship between the predictor variable (Microbial genera) and the predicted variable (Main metabolic parameters).

The indirect effect was calculated as the product of coefficients a and b, while the total effect was calculated as the sum of c and (a × b)

This approach allowed exploration of how the mediating variable (Immunological parameters) might partially explain the relationship between the predictor variables (Microbial genera) and the predicted variables (Main metabolic parameters), providing insight into the underlying mechanism of the influence of microbial genera on metabolic parameters. The Lavaan R package was used for the analysis (**36**).

## Results

### 3.1 Effect of Diets on Body Weight

A progressive increase in the mice’s body weight was observed when assessing the impact of diets along the weeks. It is noteworthy that the obesogenic diets (D2 and D3), induced a significant weight gain 4 weeks after initiating differential feeding, promoting the development of diet-induced obesity (DIO) in 100% of the mice (p-value = 0.0011); the significant difference in weights was maintained throughout the weeks when compared with the control diet (Week 12: p-value = 0.0099, Week 24: p-value = 0.0166) (Figure A-D). On the other hand, when examining the dynamics of weight gain, measured as the delta weight among different weeks, a significant difference was observed among groups; mice fed with D2 and D3 compared to D1 (p-value < 0.0001 and 0.0073), and D2 compared to D3 showed a greater delta weight (p-value = 0.0163) in week 4. No significant changes were observed when evaluating delta weights between weeks 12-4 and weeks 24-12 (ANOVA p-value = 0.136 and 0.799 respectively) (Figures 1E, 1F and 1G). Importantly, there were no significant differences in food intake among groups.

**Figure 1.**
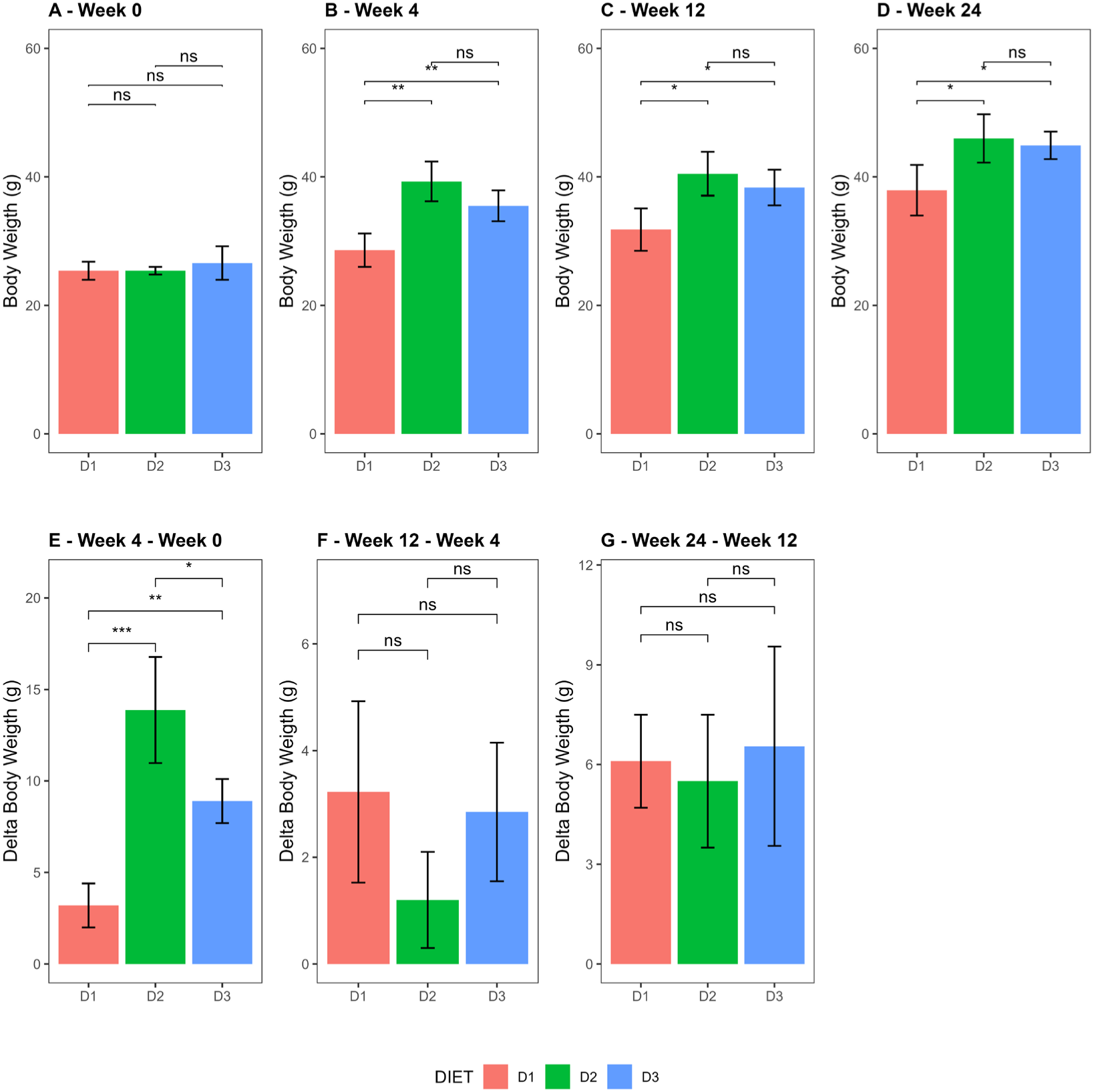
Body weight comparison. (**A**–**D**) Bar plots representing the body weight of each group, for weeks 0 (**A**), 4 (**B**), 12 (**C**), and 24 (**D**). (**E**–**G**) Bar plots representing the delta weights between different weeks at weeks 0–4 (**E**), weeks 4–12 (**F**), and weeks 12–24 (**G**). Group D1 (pink), Group D2 (green) and Group D3 (blue). Data are shown as the mean ± SD of four mice per group. (*: p ≤ 0.05, **: p ≤ 0.01, ***: p ≤ 0.001, ns: no significant).

### 3.2. Gut Microbiota Analysis

#### 3.2.1. Alpha Diversity

Alpha diversity was analyzed using observed ASV, Shannon and Simpson indexes, after 24 weeks of dietary modification. Although no significant differences were found in the observed ASVs among the groups, a significant increase in the Shannon index was observed in Group D2 compared to Groups D1 and D3 (p-value = 0.0286 in both cases). Instead, the Simpson index was increased only when comparing D2 with the control group (p-value = 0.0286). Although a trend to decreased values in all parameters analyzed was observed in Group D3 compared to D1, the differences did not reach statistical significance (p-value = 0.6860, 0.3430, and 0.6860, respectively) Figure 2.

**Figure 2.**
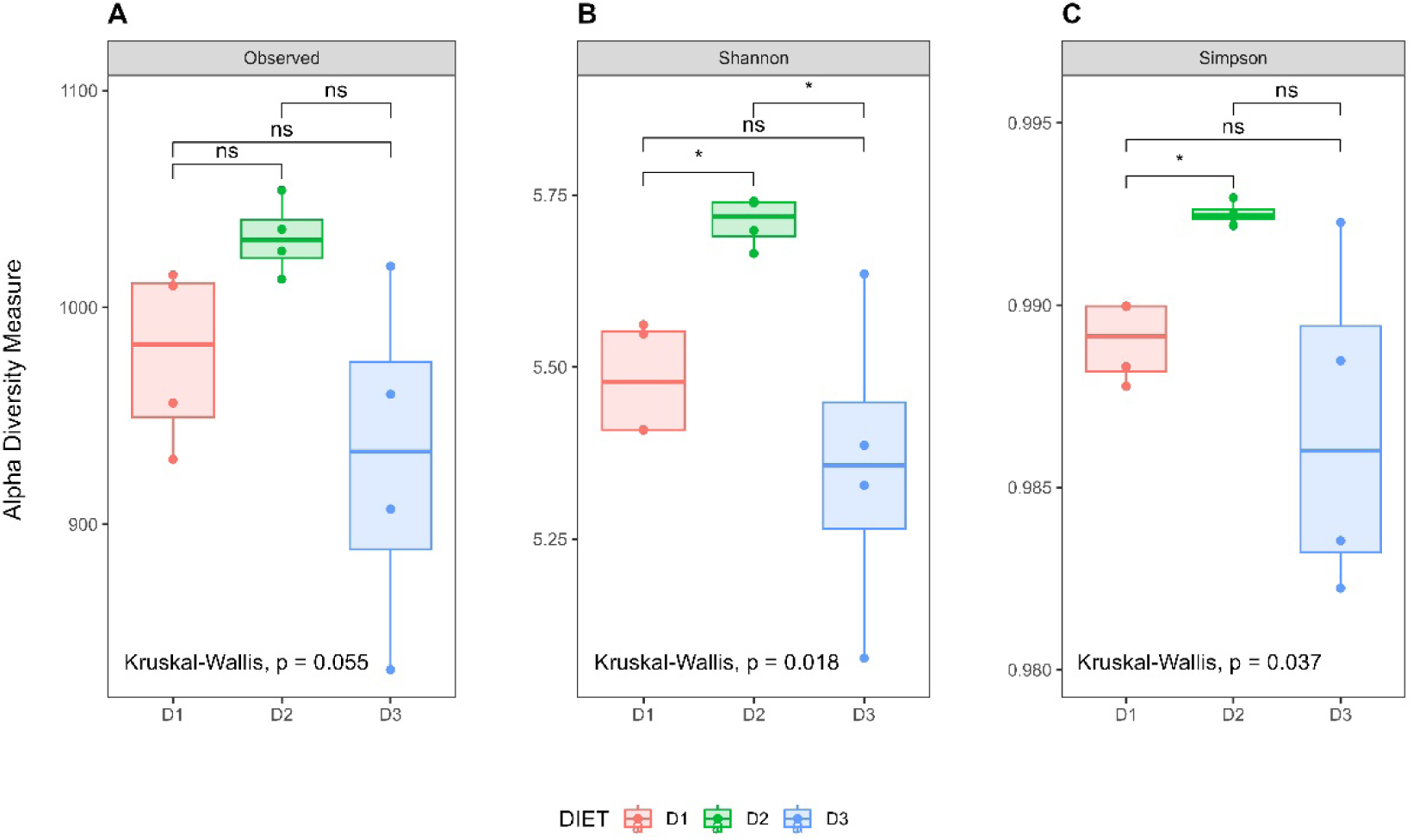
Analysis of alpha diversity among diet groups. Box plots showing observed ASV index (**A**), Shannon index (**B**), and Simpson index (**C**) of each group, Group D1 (pink), Group D2 (green) and Group D3 (blue). The solid black lines indicate the medians, and the lower and upper bounds of the box represent the 25 and 75% quartiles. Outliers are indicated as circles outside the boxes and represent samples falling outside the 10 and 90% quartiles. (*: p ≤ 0.05, ns = no significant).

#### 3.2.2 Beta Diversity Analysis

Beta diversity analysis using Principal Component Analysis (PCA) revealed clear segregation among the different groups, both visually and statistically, suggesting possible significant variations in microbiota composition (Adonis test R^2^: 0.4150, p-value = 0.001, ANOSIM R^2^ = 0.9583, p-value = 0.001). Groups D2 and D3, which received a medium-fat content diet, exhibit greater similarity to each other compared to the control group D1 (Figure 3A). Results obtained with Unifrac analysis in both weighted and unweighted approaches were consistent with PCA. Visually, some group overlap was observed in weighted, contrasting with unweighted Unifrac. However, greater discriminative power was identified in weighted Unifrac (ANOSIM R^2^: 0.6273, p-value = 0.001) compared to unweighted Unifrac (ANOSIM R^2^: 0.3426, p-value = 0.003), indicating that variations in microbiota composition among groups are more pronounced when considering the relative abundances of taxa (Figures 3B and 3C).

**Figure 3.**
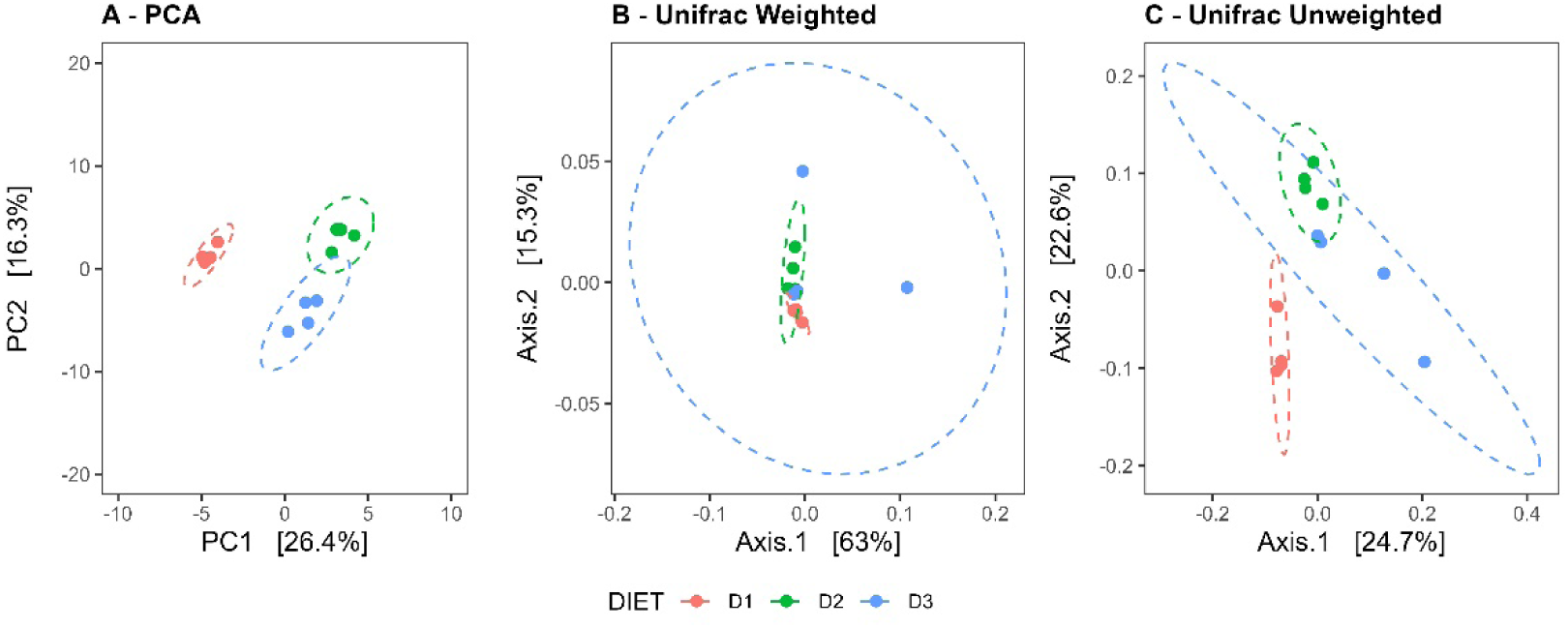
Beta diversity analysis. (**A**) Comparison of the microbiota profiles using Principal Component Analysis (PCA). The first two principal components, PC1 and PC2, were plotted (Adonis test R2: 0.4150, p-value = 0.001, ANOSIM R2 = 0.9583, p-value = 0.001). Comparisons based on the weighted (**B**) and unweighted (**C**)UniFrac distances respectively (ANOSIM R2: 0.6273, p-value = 0.001; and ANOSIM R2: 0.3426, p-value = 0.003). Group D1 (pink), Group D2 (green) and Group D3 (blue)

### 3.3 LEfSe Differential Analysis

To identify taxa enriched in the different groups according to the diet received, Linear Discriminant Analysis (LDA) together with effect size measurements (LEfSe) was used. LDA score of genera enriched in each group in the global analysis is shown in Figure 4. Pairwise comparisons to obtain a detailed understanding of the effects of diet on gut microbiota composition is also presented (Figure 5A, 5B and 5C).

**Figure 4.**
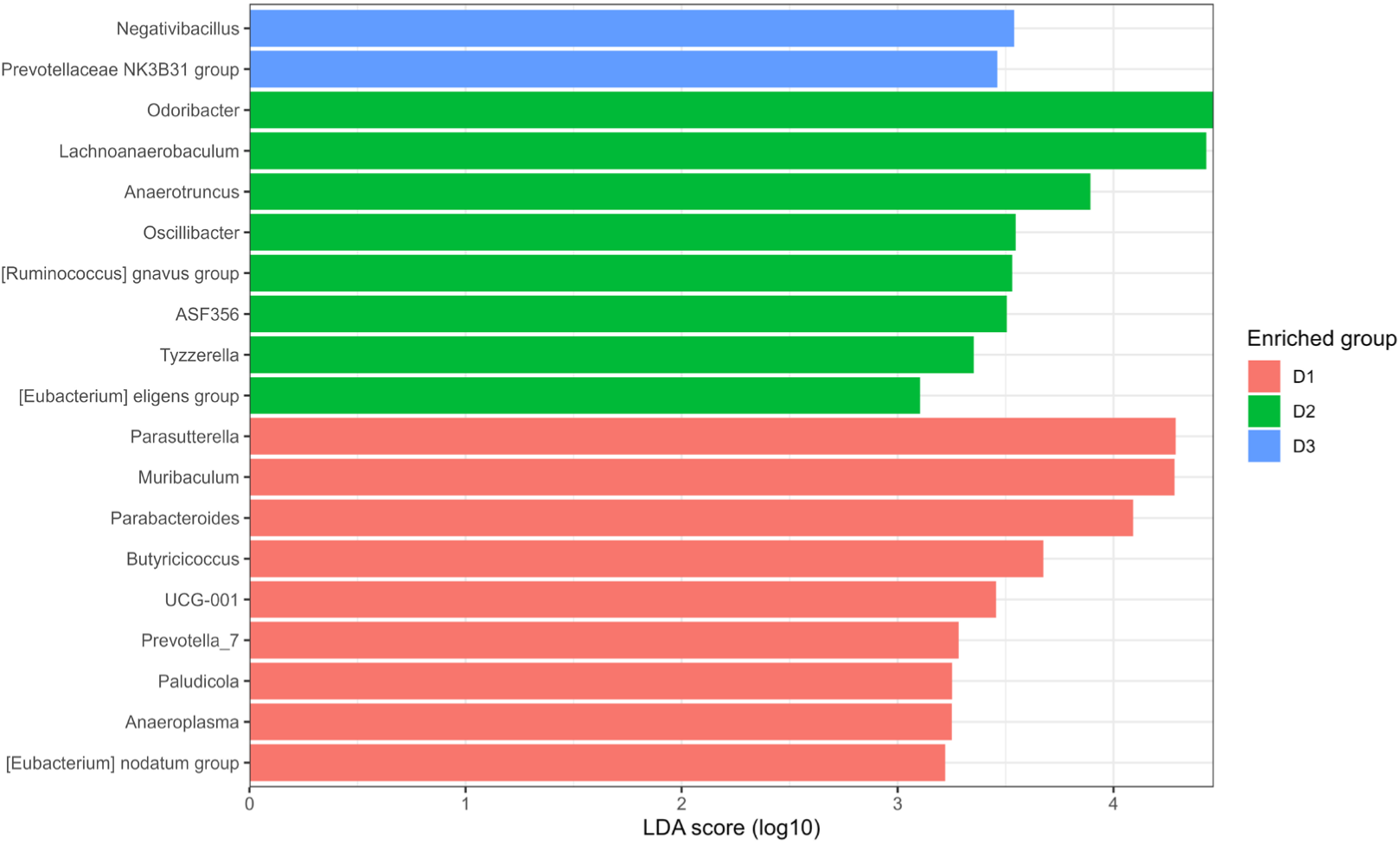
Linear discriminant analysis (LDA) coupled with effect size measurements (LEfSe) for global analysis. Enriched taxa are represented by a positive LDA score: Group D1 (pink), Group D2 (green) and Group D3 (blue). Scores > 3 and *p*-value < 0.05 obtained by the Kruskal–Wallis test were considered significant.

**Figure 5.**
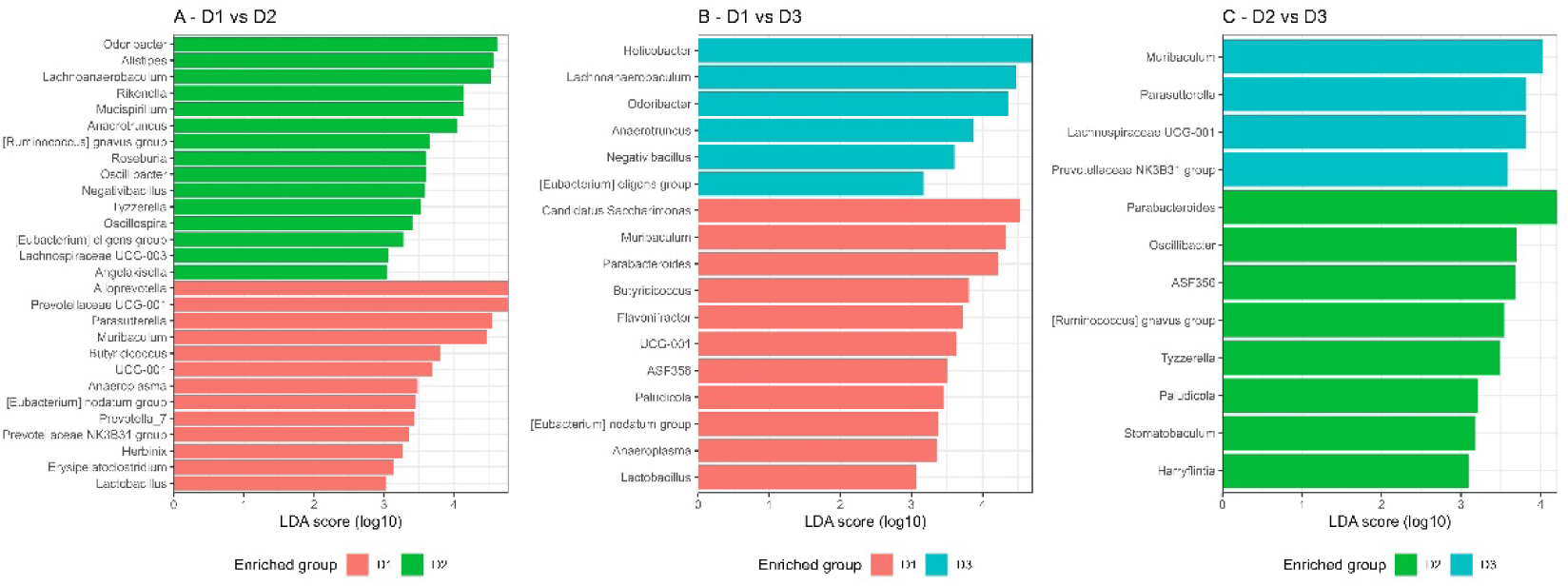
Linear discriminant analysis (LDA) coupled with effect size measurements (LEfSe) for Pairwise comparisons **(A)** D1 vs D2, (**B**) D1 vs D3 and (**C**) D2 vs D3. Enriched taxa are represented by a positive LDA score: Group D1 (pink), Group D2 (green) and Group D3 (blue). Scores > 3 and *p*-value < 0.05 obtained by the Kruskal–Wallis test were considered significant.

When comparing both D2 and D3 against D1, we observed that the genera *Lachnoanaerobaculum*, *Odoribacter*, *Anaerotruncus*, *Negativibacillus*, *[Eubacterium] eligens group*, were enriched in both fat diets fed groups (D2 and D3). When comparing D2 vs D3, we observed that D3 is mainly enriched with genera: *Muribaculum*, *Parasutterella*, and *Prevotellaceae NK3B31*, found as well in D1 when D1 was compared to D2. At the same time, D2 was enriched in certain genera, *Parabacteroides, ASD356*, and *Paludicola* present in D3, when D3 was compared with D1, as well as *Tyzzerella, [Ruminococcus] gnavus group* and *Oscillibact*er.

### 3.4 Metabolic Status and Immune Cell Populations Profiling in VAT

The assessment of the metabolic status demonstrates that leptin levels in Groups D2 and D3 were higher than those in Group D1 (Figure 6A), according to body weight differences shown by the different groups of mice (D1 vs D2: p-value = 0.0025, D1 vs D3: p-value = 0.0016 and D2 vs D3: p-value = 0.999, respectively). In contrast, adiponectin levels decreased in Groups D2 and D3 compared to the control diet (Figure 6B). Although these differences did not reach statistical significance (ANOVA p-value = 0.206), it is important to note that the values in mice of Group D3 were lower than in Group D2. Additionally, a significant increase total cholesterol levels were found in Group D2 compared to Group D1 (p-value = 0.0063 respectively) (Figure 6C). No other statistically significant changes were observed in the remaining parameters evaluated, namely glucose, triglycerides, total proteins, albumin, ALT, and AST (Figures 6D, 6E, 6F, 6G, 6H and 6I).

**Figure 6.**
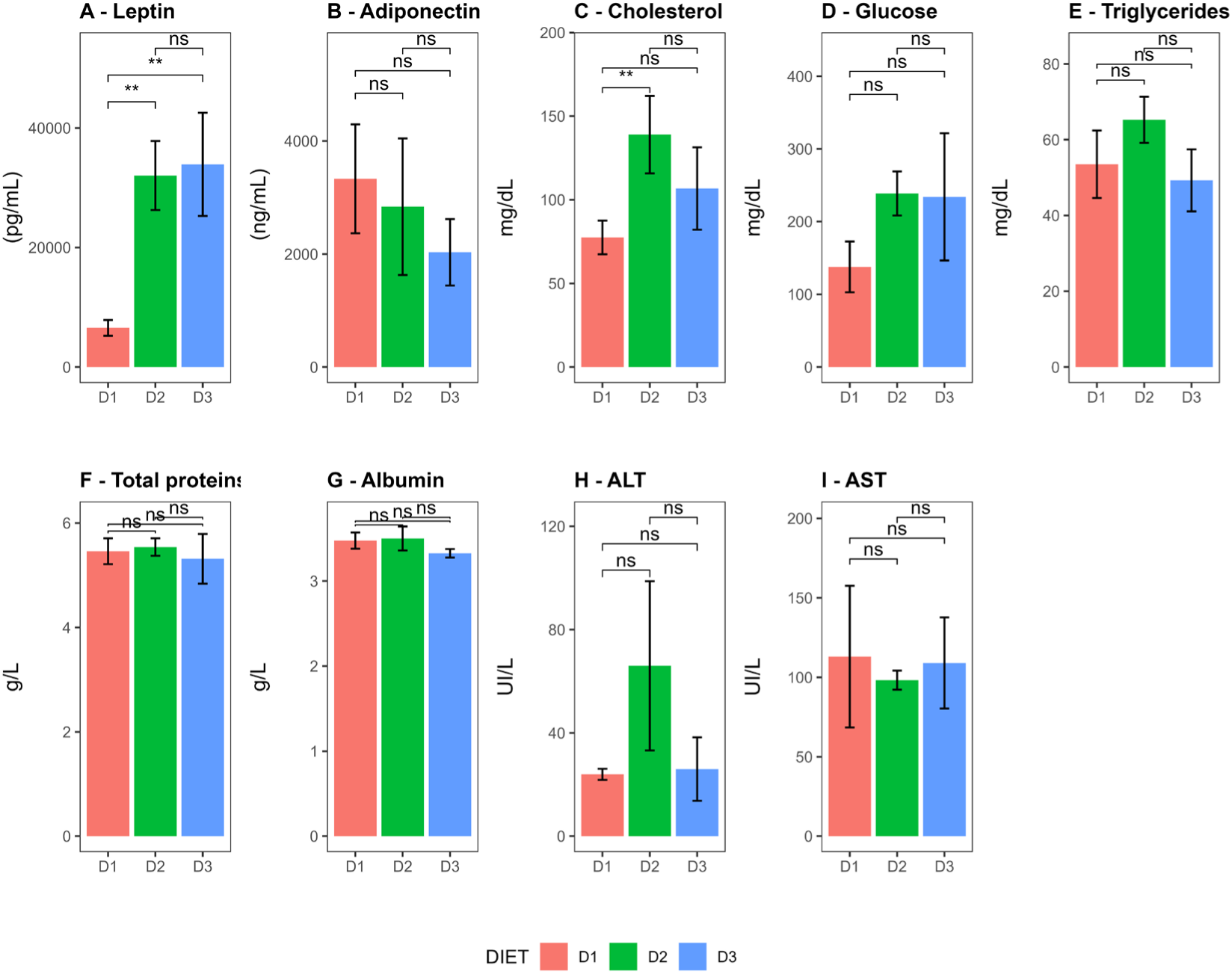
Blood metabolic parameter comparison among groups. (**A**–**I**) Concentrations of blood metabolic parameters, namely leptin, adiponectin, total cholesterol, glucose, triglycerides, total proteins, albumin, alanine aminotransferase (ALT) and aspartate aminotransferase (AST). Bar plot representing the parameters evaluated for each group, Group D1 (pink), Group D2 (green) and Group D3 (blue)., at week 24 post-starting differential feeding. Data are shown as the mean ± SD of four mice per group. (*: p ≤ 0.05, **: *p* ≤ 0.01 and ns = no significant).

The characterization of immune cells in the VAT showed that mice fed with the diet D2 had a significantly higher proportion of regulatory T cells (CD45+ CD4+ FoxP3+) compared to group D1 (p-value = 0.0002). The same was observed when comparing D3 vs D1, (p-value = 0.0051) (Figure 7A). When analyzing the expression of IL-10, both: the percentage of IL-10+ cells and the Mean Fluorescence Intensity (MFI) of IL-10 in Tregs cells the results showed lower values in the groups fed with a fatty diet compared to the control group (Figure 7B and 7C). The statistical significance for percentage of IL-10+ cells (CD45+ CD4+ FOXP3+ IL-10+) was (D1 vs D2: p-value = 0.0341, D3 vs D1: p-value = 0.0988, and D2 vs D3: p-value = 1.0000) and for the MFI was (0.029) both for the comparison D1 vs D2 and D1 vs D3 and (0.486) por D2 vs D3. Additionally, the proportions of myeloid cells (CD45+, CD11b+), macrophages (CD45+, CD11b+, F4/80+), and CD4 lymphocytes (CD45+ CD4+) were also similar across all groups (Figure 7D, 7E and 7F).

**Figure 7.**
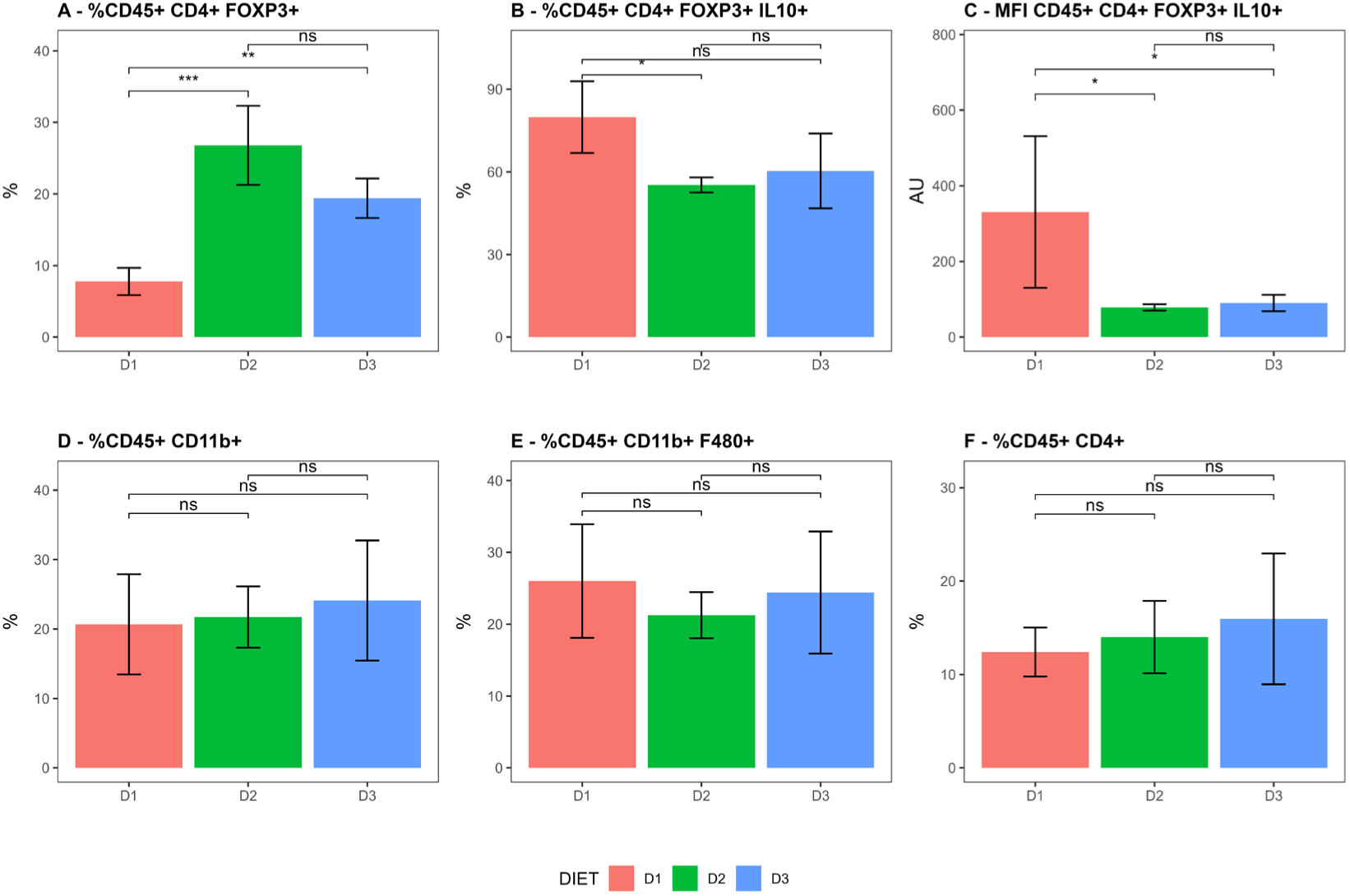
Comparison of immune cell population profiles among diet Groups. Bar plots representing: (**A**) % CD45+ CD4+ FOXP3+ cells, (**B**) % CD45+ CD4+ FOXP3+ IL-10+ cells, (**C**) MFI for CD45+ CD4+ FOXP3+ cells, (**D**) % CD45+ CD11b+ cells, (**E**) % CD45+ CD11b+ F4/80+ cells and (**F**) % CD45+ CD4+ cells for each group, Group D1 (pink), Group D2 (green) and Group D3 (blue)., at week 24 post-starting differential feeding. Data are shown as the mean ± SD of four mice per group. (*: p ≤ 0.05, **: *p* ≤ 0.01, ***: *p* ≤ 0.001 and ns = no significant). MFI: mean fluorescence intensity, CD: cluster of differentiation, FOXP3: forkhead box P3, IL-10: interleukin 10, AU: arbitrary units.

**Figure 8.**
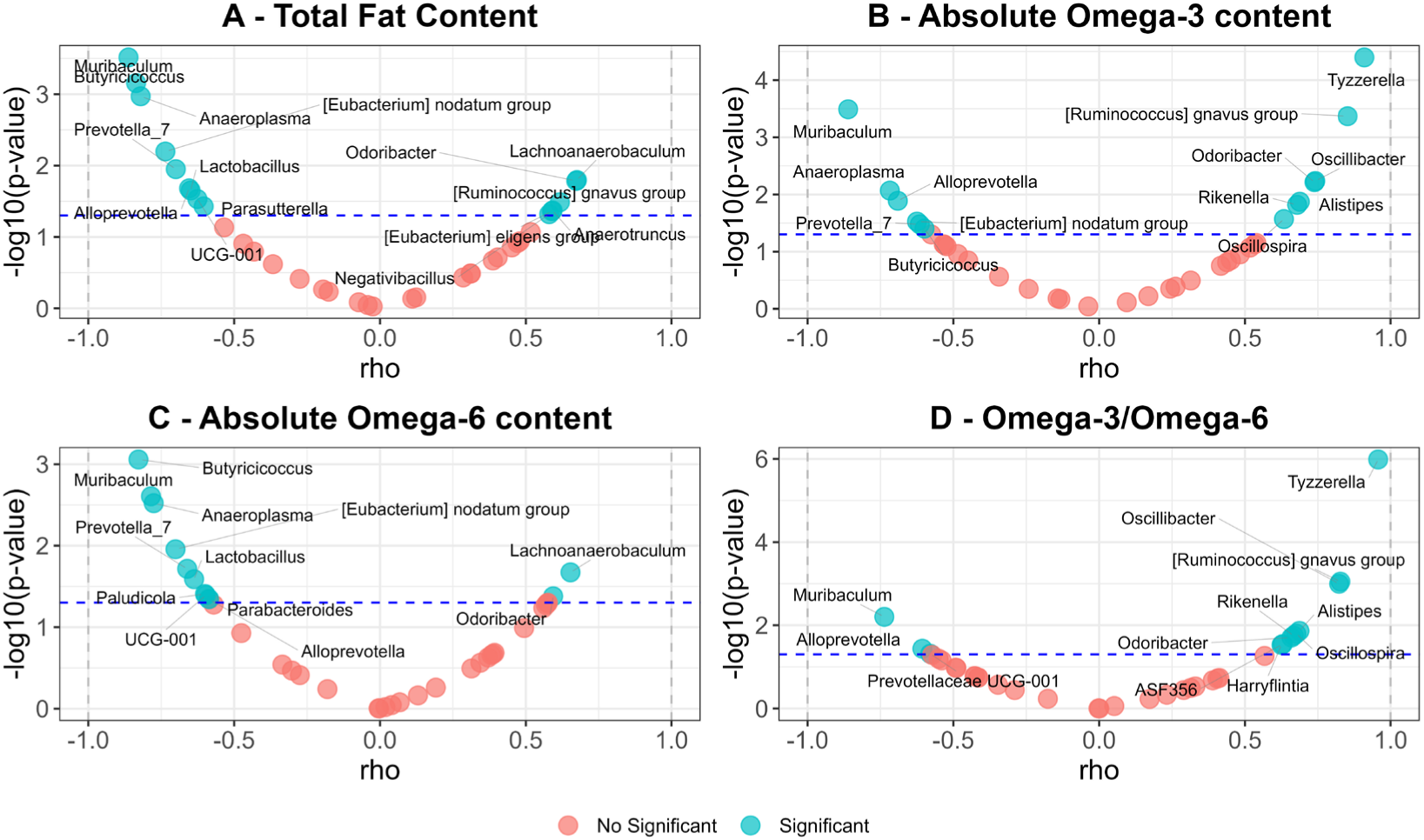
illustrates the correlation analysis between the enriched bacterial genera and the dietary fat content; (**A**): Total fat content, (**B**): Absolute Omega 3 content, (**C**): Absolute Omega 6 content and (**D**): Omega 3 / Omega 6 ratio.. The points represent the correlation (rho) on the x-axis and the negative base 10 logarithm of the p-value (-log10(p-value)) on the y-axis. Points are colored based on whether the p-value is significant (blue) or not (red). Dashed lines indicate the significance threshold (alpha) (0.05). Additionally, significant variables are labeled with repulsive text to avoid overlaps.

### 3.4 Correlation Analysis

Volcano plots in Figure 8 show the correlation analysis between the GM bacterial genera enriched in each group and the dietary fat content. Tyzzerella*, Ruminococcus gnavus group, Oscillibacter Alistipes, Rikenella, Oscillospira,* and *Odoribacter* correlate positively and significantly with the absolute content of Omega-3 in the diet. These 7 genera, along with *ASF356* and *Harryflintia*, also correlated positively with the Omega-3/Omega-6 ratio. In contrast, *Muribaculum, Anaeroplasma, Alloprevotella, Prevotella_7, Butyricicoccus*, and *[Eubacterium] nodatum group* negatively correlated with total fat, absolute Omega-3, and absolute Omega-6 content.

The same analysis was done to correlate the immune cells present in VAT with bacterial genera showing differential abundance. As shown in Figure 9, the relative abundance of *Negativibacillus, [Ruminococcus] gnavus group, Alistipes, Odoribacter,* and *Tyzzerella* correlated significantly and positively with the abundance of Tregs. Conversely, *Alloprevotella, Prevotellaceae UCG-001, Parasutterella, Muribaculum, Prevotella_7, Anaeroplasma, UCG-001, Butyricicoccus,* and *HT002* did so negatively. This pattern is very similar to that observed when the absolute amount of Omega-3 in the diet was compared.

**Figure 9.**
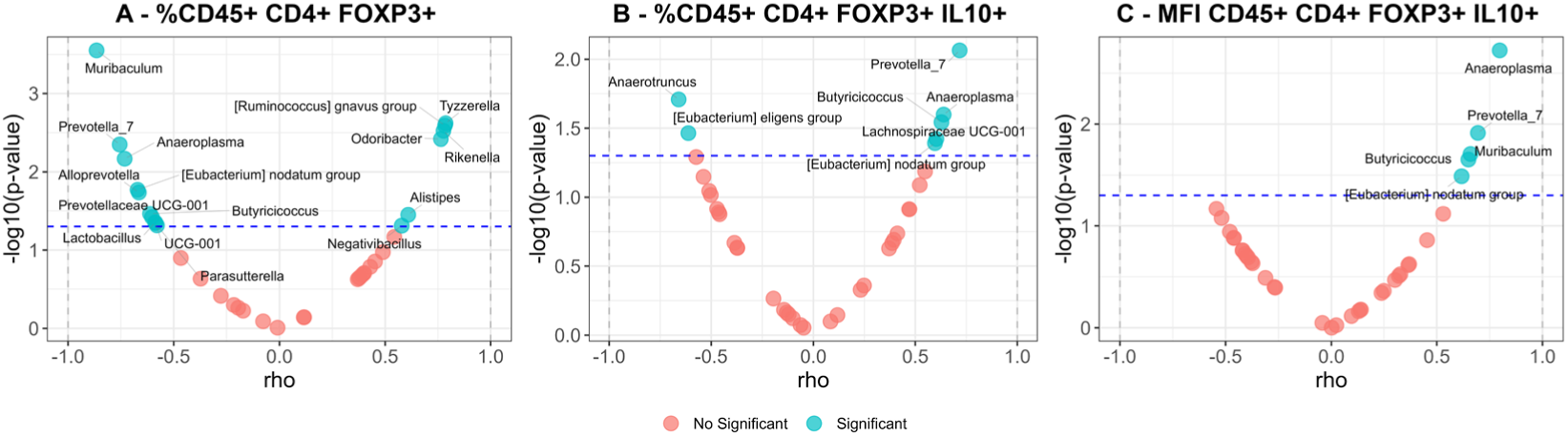
illustrates the correlation analysis between the enriched bacterial genera and immunological parameters in VAT: (**A**): %CD45+ CD4+ FOXP3+ cells, (**B**): %CD45+ CD4+ FOXP3+ IL-10+ cells and (**C**): MFI of CD45+ CD4+ FOXP3+ cells. The points represent the correlation (rho) on the x-axis and the negative base 10 logarithm of the p-value (- log10(p-value)) on the y-axis. Points are colored based on whether the p-value is significant (blue) or not (red). Dashed lines indicate the significance threshold (alpha) (0.05). Additionally, significant variables are labeled with repulsive text to avoid overlaps. MFI: *mean fluorescence intensity,* CD: cluster of differentiation. FOXP3: forkhead box P3, IL-10: interleukin 10

The percentage of CD4+ Tregs cells (CD45+ CD4+ FOXP3+) expressing IL-10 correlated negatively with the relative abundance of *Anaerotruncus* and *Eubacterium eligens group*, and positively with *Prevotella_7, Butyricicoccus, Anaeroplasma, Laschnospiraceae UCG-001*, and *Eubacterium nodatum group*. These five genera, along with *Muribaculum*, also correlated positively with the MFI of this regulatory population.

When metabolic markers and Treg cells profile were correlated, (Figure 10), the percentage of CD4+ Tregs (CD45+ CD4+ FOXP3+) correlated positively with levels of glucose, leptin, ALT, and cholesterol. However, these correlations became negative for glucose and cholesterol when considering the percentage of cells expressing IL-10 molecule. Similarly, when evaluating the MFI of IL-10 positive cells, negative correlations were found with glucose and leptin levels, with a positive correlation for adiponectin.

**Figure 10.**
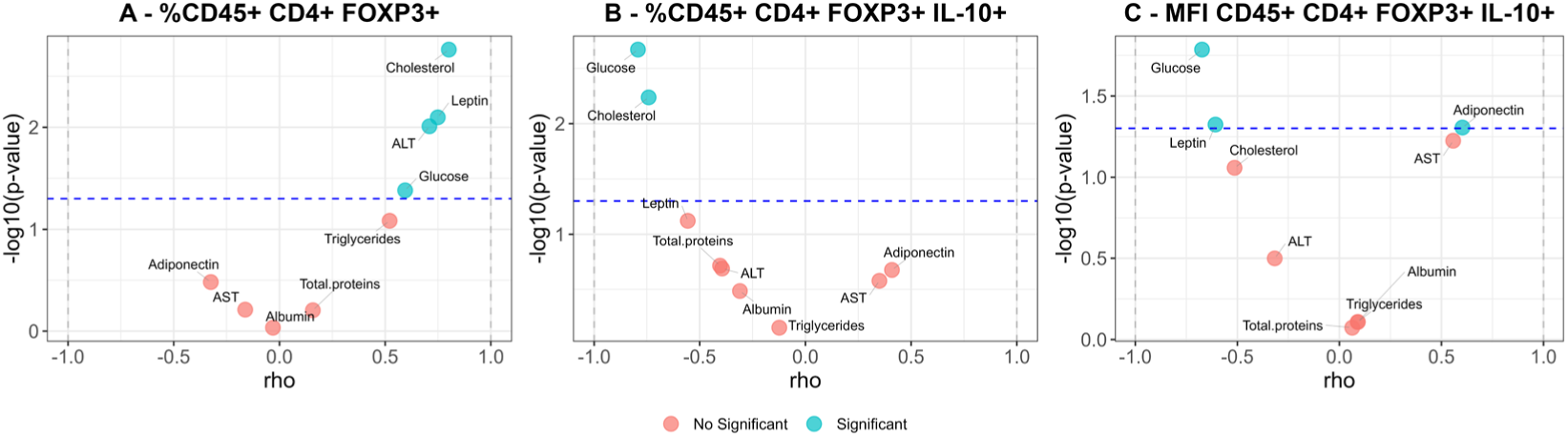
illustrates the correlation analysis between metabolic and immunological parameters in VAT: (**A**): %CD45+ CD4+ FOXP3+ cells, (**B**): %CD45+ CD4+ FOXP3+ IL-10+ cells and (**C**): MFI of CD45+ CD4+ FOXP3+ cells. The points represent the correlation (rho) on the x-axis and the negative base 10 logarithm of the p-value (-log10(p-value)) on the y-axis. Points are colored based on whether the p-value is significant (blue) or not (red). Dashed lines indicate the significance threshold (alpha) (0.05). Additionally, significant variables are labeled with repulsive text to avoid overlaps. MFI: mean fluorescence intensity, CD: cluster of differentiation. FOXP3: forkhead box P3, IL-10: interleukin 10, ALT: Alanine Aminotransferase

Figure 11 show correlation between metabolic markers with the enriched bacterial taxa, a similar profile to the found in the correlation analysis between bacterial taxa and dietary fat content was found.

To determine if the bacterial genera enriched in the groups were modifying metabolic parameters through immunological modulation of cells in VAT, a mediation analysis was conducted. The parameters considered were Glucose, Cholesterol, Leptin, Adiponectin, % of CD45+ CD4+ FOXP3+ cells, % of CD45+ CD4+ FOXP3+ IL10+ cells and MFI of CD45+ CD4+ FOXP3+ IL10+ cells. Figure 12 shows a scheme representing the mediation models and regression pathways, while Supplementary Table S6 presents the values obtained. Figure 13 displays the standardized estimates for each of the mediations performed. Only those mediations showing a significant indirect effect (p-value <0.05) are plotted.

**Figure 11.**
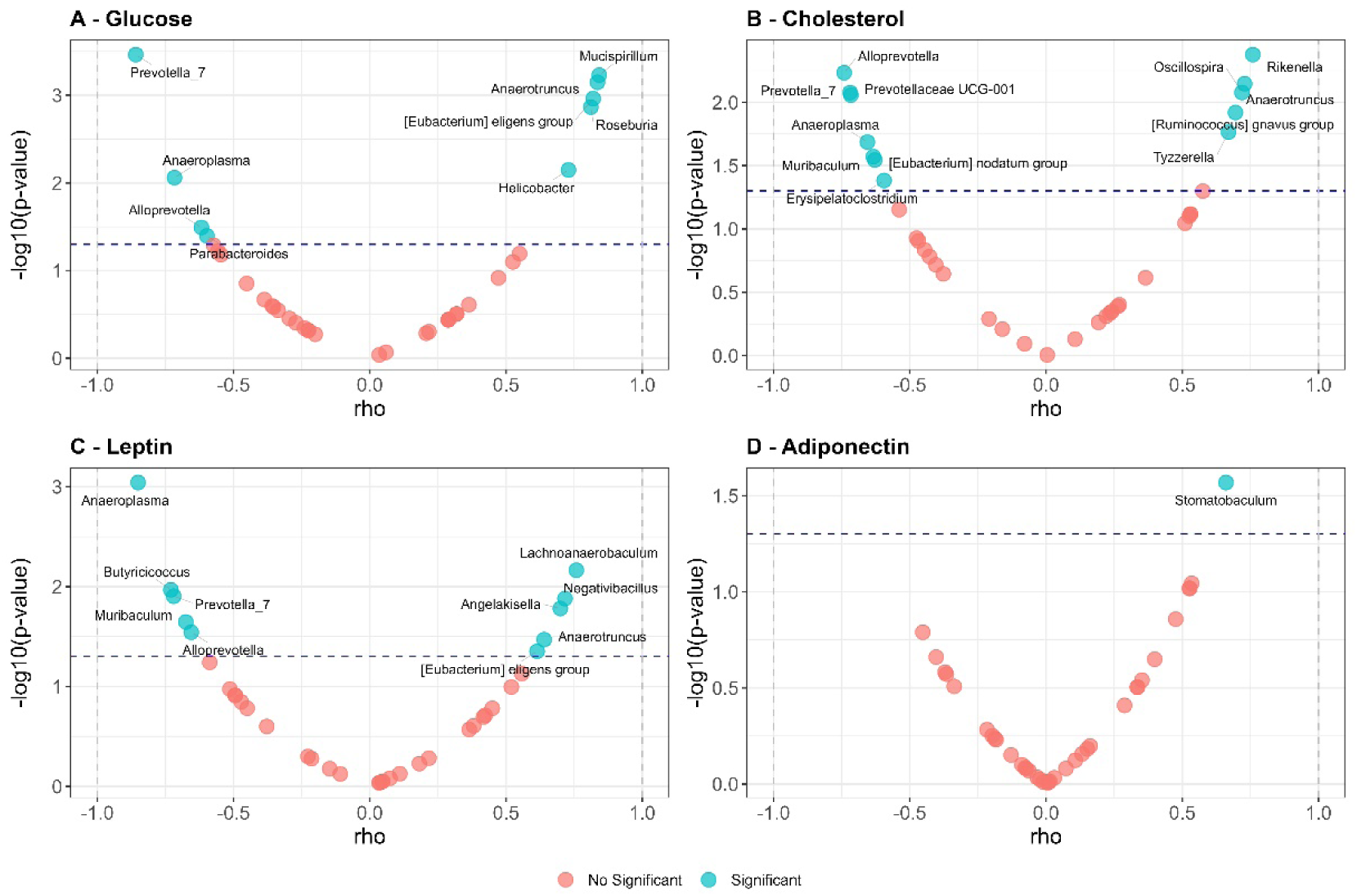
illustrates the correlation analysis between the enriched bacterial genera and main metabolic parameters; (**A**): Glucose, (**B**): Cholesterol, (**C**): Leptin and (**D**): Adiponectin. The points represent the correlation (rho) on the x-axis and the negative base 10 logarithm of the p-value (-log10(p-value)) on the y-axis. Points are colored based on whether the p-value is significant (blue) or not (red). Dashed lines indicate the significance threshold (alpha) (0.05). Additionally, significant variables are labeled with repulsive text to avoid overlaps.

**Figure 12.**
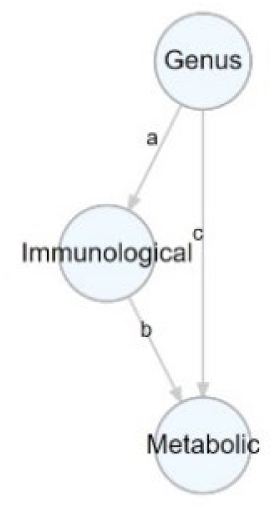
shows the mediation model used: The nodes represent; Genus: (Enrichment bacterial taxa), Immunological: (Immunological parameters) and Metabolic: (Metabolic parameters). Edges labeled ’a’, ’b’ and ’c’ connect the nodes and demonstrate the relationships within the model: a: represents the relationship between the predictor variable (microbial genus) and the mediator variable (immunological parameter). b: represents the relationship between the mediating variable (immunological parameter) and the predicted variable (Main metabolic parameter). c: represents the direct relationship between the predictor variable (microbial genus) and the predicted variable (Main metabolic parameter).

**Figure 13.**
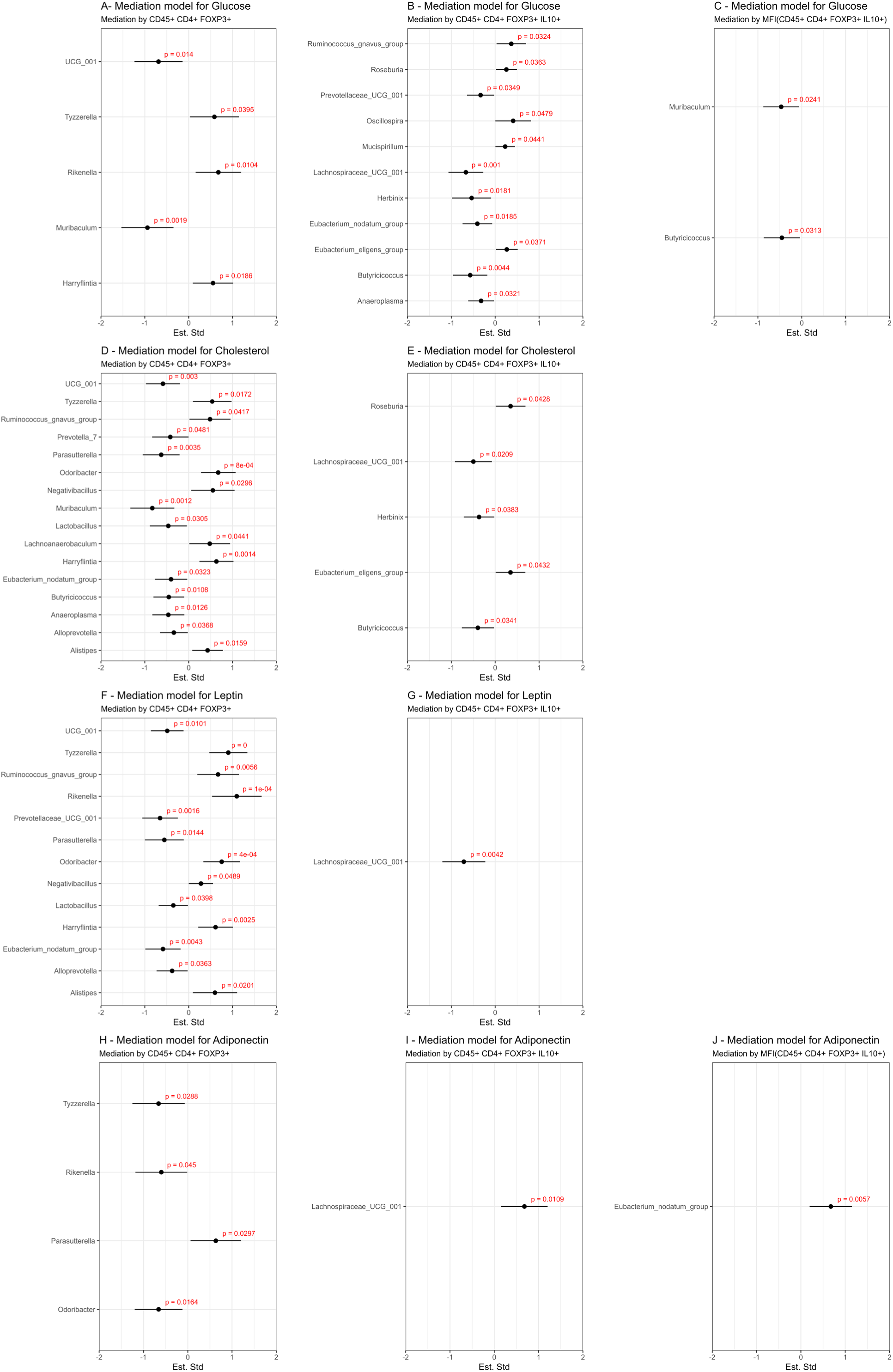
displays the standardized estimates for each of the conducted mediations. Only those mediations showing a significant indirect effect (p-value < 0.05) are graphically represented. (**A**): Glucose values mediated by % of CD45+ CD4+ FOXP3+ cells. (**B**): Glucose values mediated by % of CD45+ CD4+ FOXP3+ IL-10+ cells. (**C**): Glucose values mediated by the MFI of CD45+ CD4+ FOXP3+ IL-10+ cells. (**D**): Cholesterol values mediated by % of CD45+ CD4+ FOXP3+ cells. (**E**): Cholesterol values mediated by % of CD45+ CD4+ FOXP3+ IL-10+ cells. (**F**): Leptin values mediated by % of CD45+ CD4+ FOXP3+ cells. (**G**): Leptin values mediated by % of CD45+ CD4+ FOXP3+ IL-10+ cells. (**H**): Adiponectin values mediated by % of CD45+ CD4+ FOXP3+ cells. (**I**): Adiponectin values mediated by % of CD45+ CD4+ FOXP3+ IL-10+ cells. (**J**): Adiponectin values mediated by the MFI of CD45+ CD4+ FOXP3+ IL-10+ cells. The models for cholesterol values mediated by the MFI of CD45+ CD4+ FOXP3+ IL-10+ cells, and leptin values mediated by the MFI of CD45+ CD4+ FOXP3+ IL-10+ cells, are not represented as they did not yield significant indirect effect values. Each point on the graph indicates the standardized effect in the mediation for the enriched bacterial genus. The error bars show the confidence intervals for these standardized effects. The p-values for each effect are also indicated. MFI: mean fluorescence intensity, CD: cluster of differentiation. FOXP3: forkhead box P3, IL-10: interleukin 10,

## 4. Discussion

Dietary fat plays an important role in modulating the composition of the GM. Certain fatty acids escape digestion in the small intestine and reach the colon, where they can be metabolized by the GM. Although numerous studies support the evidence that a high fat diet (HFD) can alter the composition of the GM and have negative health consequences (**12,13,37,38**), there still exists a gap in understanding how different dietary lipids may influence specific members of the microbiota and generate both beneficial and detrimental health consequences. Our previous study in a murine model, demonstrated that sustained administration of Omega-3 fatty acids in the diet induces specific changes in the composition of the GM, modulating the immune response in VAT and maintaining adequate metabolism despite the development of obesity (**39**). In the present study, we aimed to determine if changes in the GM induced by constant administration of Omega-3 are associated with the induction of Tregs in VAT and if this effect is dose-dependent.

Initially, we observed that diets with medium-fat content have an obesogenic effect over a period of 4 weeks, regardless of the levels of Omega-3 supplied, which is expected since the energy intake is similar for each of them. Comparable effects have been observed by LeMieux et al., when comparing HFD vs. HFD plus added eicosapentaenoic acid (EPA) (**40**). However, after the development of obesity, we noticed a stabilization of body weight in the groups that received the experimental diets, which ,as suggested in previous studies, could be attributed to the regulation of metabolism induced by the Omega-3 fatty acids, (**39,41,42**)

Our results indicate that dietary modification did not significantly affect the diversity of microbial species present in terms of ASV observed but did alter the uniformity of their relative abundance distribution. Furthermore, an increase in the Simpson index was observed in the group D2 receiving a higher amount of Omega-3, compared to the control group. Since the Simpson index is a measure of diversity assigning more weight to dominant species our finding suggests that dietary modification led to a shift in the composition of the microbiota towards less dominant species in Group D2. Although no significant differences in Alpha diversity Indexes were observed when comparing D1 vs. D3 a trend towards lower values in D3 was noted. Multiple studies have highlighted a relationship between obesity and a reduction in microbial diversity (**43,44**). However, it is important to note that while individuals with less diverse GM tend to be more obese, this low diversity is not a common characteristic in most obese populations (**45**), suggesting that low diversity may be relevant only in certain subgroups of individuals with obesity (**46**). Furthermore, consistent with our results, it has been reported that HFD tend to reduce microbial diversity, while supplementation with Omega-3 may promote its increase (**47**). We hypothesize that the positive effects observed on microbial diversity in our model may be related to the amount of Omega-3 provided in the diet or a higher Omega-3/Omega-6 ratio. As our results suggest, only those mice receiving a higher amount of Omega-3 showed these changes.

Beta diversity analysis of the GM revealed that groups receiving a fat diet exhibited greater similarity to each other compared to the control group. This pattern suggests a specific influence of the dietary fats on the structure of the GM. In both groups, an enrichment of certain microorganisms associated with the production of short-chain fatty acids (SCFA) such as *Lachnoanaerobaculum, Odoribacter, Anaerotruncus, Negativibacillus*, and *[Eubacterium] eligens group* was observed. It is noteworthy that previous studies by Watson et al., have reported an increase in the abundance of bacterial producing butyrate after supplementation with Omega-3 fatty acids (**48**). On the other hand, mice in the control group showed as well enrichment in certain genera associated with SCFA production, such as *Eubacterium nodatum group, Parasutterella, Parabacteroides, Butyricicoccus*, and *Prevotella_7*. It is very well known that SCFA can be used for the biosynthesis of lipids, cholesterol, and proteins, and act as signaling molecules for host tissues and organs, mediating effects related to oxidative stress, immune response, insulin resistance, and inflammatory responses (**49**). Although high SCFA production is commonly associated with a healthy GM and the presence of microorganisms related to reduction of obesity, it is important to note that increased SCFA production has also been proposed as a mechanism linked to the development of obesity (**50**). Therefore, it is crucial to delve into the molecular study of the metabolic pathways underlying the microbial communities present here to understand how they can modulate the individual’s energy balance. Additionally, *Muribaculum*, a genus associated with markers of intestinal barrier function (**51**) was found to be enriched in Group D1 compared to Groups D2 and D3; this was previously identified as a microbial signature for the chow diet (**39**). Similar results were presented by Huang et al., when studying mice fed a HFD compared to Chow Diet, (**52**). It is important to highlight that among the genera with higher abundance in mice receiving a fat diet are *Odoribacter, Alistipes, Lachnoanaerobaculum, Mucispirillum, Anaerotruncus, Oscillibacter, Roseburia*, and *Helicobacter*, which are part of the core intestinal bacteria identified in healthy mice by Wang et al. (**53**). This finding is significant, since variations in the distribution of these taxa could be linked to the immunometabolic consequences associated with fat intake and Omega-3 supplementation.

Obesity development is characterized by AT accumulation triggering dysregulation of adipokine production with leptin and adiponectin, significantly contributing to obesity-related disease onset (**54,55**). In our experimental model, we found that leptin a key hormone in food intake, body mass regulation, and various physiological functions, presented higher levels in Groups D2 and D3 compared to the control group; this is in agreement with the differences in body weight among groups (**56**). On the other hand, adiponectin, a hormone with a broad spectrum of influences on metabolic processes, anti-inflammatory properties and insulin-sensitizing capacity (**57**) showed no significant differences in levels among the groups studied. However, it is important to note that increased visceral fat in overweight or obese individuals is associated with decreased total adiponectin levels (**58**). This reduction could contribute to insulin resistance and inflammation associated with obesity and other metabolic conditions. It is noteworthy that while we observed a trend towards lower adiponectin levels in obese mice, this trend is even more pronounced in those mice receiving lower Omega-3 fatty acid. This finding is significant as circulating adiponectin levels reflect its metabolic impact on adipocytes and adipose tissue (**59**). Additionally, the omega-6/Omega-3 fatty acids ratio inversely correlates with adiponectin levels, suggesting a fundamental role of this ratio in metabolic homeostasis and cardiovascular health (**60**). These findings may partly explain the preservation of an adequate metabolic profile in our model, without alterations in the main metabolites analyzed such as glucose, triglycerides, AST and ALT. On the other hand, the elevated total cholesterol levels observed in Group D2 may be attributed to a significant increase in HDL cholesterol levels, as demonstrated in our previous publication (**39**). Lastly, numerous studies support the notion that adequate adiponectin levels are associated with good metabolic management (**61–63**)

It is very well documented that obesity induces inflammation in VAT characterized by an imbalance between local proinflammatory T cells and Tregs, contributing to metabolic abnormalities associated with this condition. When we studied the immune cells in VAT, our results revealed that mice fed with fat diet exhibited a significantly higher proportion of regulatory T cells (CD45+ CD4+ FP3+), and this proportion increased with the dose of Omega-3 provided. This finding strengthens the hypothesis that immunometabolic regulation in VAT may be mediated by the recruitment and/or induction of Tregs. Previous studies have shown that the frequency and number of VAT Tregs are drastically reduced in three mouse models of obesity, including ob/ob mice, Ay/a obese mice, and HFD-induced obese mice (**22**) and that in lean mice, Tregs can account for 40 to 80% of CD4+ T cells in VAT (**64**). This underscores the importance of Tregs in maintaining metabolic balance and immunological homeostasis. In this context, dietary supplementation with Omega-3 has been associated with the proliferation and accumulation of Tregs in various tissues, including the liver (**65**), spleen (**66**), and particularly AT in wild-type mice (**23**). In vivo studies have shown that T cell differentiation into Tregs is favored by the induction of M2 macrophages, and this effect could be mediated by the provision of Omega-3 fatty acids (**67,68**). The presence of Tregs in VAT is crucial for regulating local and systemic metabolism. Their temporary elimination led to increased activation of inflammatory genes in AT, along with insulin resistance and glucose intolerance in mice consuming a HFD, emphasizing the notion that Tregs play a significant role in curbing inflammation in AT (**69**). Additionally, SCFAs are also postulated as mediators of Treg induction in VAT (**70**). In the context of our study, Omega-3 fatty acids may have a direct effect on Treg induction, explaining their higher presence in the VAT of mice fed with fat diet and Omega-3 could influence the composition of the GM, generating a microbial structure that favors SCFA production, which in turn could mediate as well Treg induction in VAT. It is important to note that, unfortunately, in our study we did not determine circulating SCFA levels, limiting our ability to fully confirm this hypothesis. However, the accumulated evidence in the literature supports the interaction between GM composition, SCFA production and immunometabolic regulation, suggesting that this pathway is reasonable and deserves further exploration.

Additionally, it is interesting to note that, although IL-10 is recognized as one of the main cytokines associated with the regulatory function of Tregs, we observed that the groups receiving a diet rich in Omega-3 presented a lower proportion of CD45+ CD4+ FoxP3+ IL-10+ cells, with a significant difference mainly between D1 and D2. These observations are in agreement with previous research revealing significant differences in the transcriptional profiles of Treg cells in VAT between ob/ob or HFD-induced obese mice and lean mice. For example, a substantial decrease in IL-10 expression levels has been observed in obese mice. In this context, a study by Beppu et al. provides an interesting perspective, demonstrating that in mice knockout for IL-10 or its transcriptional regulator Blimp-1 in Tregs, instead of developing weight gain and insulin resistance, they were protected from these adverse effects when fed an HFD. On the other hand, high levels of IL-10 have been shown to suppress thermogenesis in adipocytes (**71**). Therefore, the suppression of Tregs IL-10 secretion led to increased adipocyte thermogenesis, resulting in reduced weight gain and improvements in insulin resistance in mice (**72**). Together, these data suggest the possibility that some of the effects of Omega-3 on Tregs and obesity may be related to their influence on Blimp-1 regulation. However, further studies are needed to validate and fully understand these effects and their implications in immunometabolic regulation.

Finally, simple mediation analyses revealed significant associations between the microbial profile and immunometabolic regulation. To remark, is the capacity of *Lachnospiraceae UCG-001* to modulate levels of leptin, glucose, and cholesterol through the stimulation of CD45+ CD4+ FOXP3+ IL10+ cells. Bacteria from the *Lachnospiraceae* family are recognized for their ability to produce SCFA; enrichment of *Lachnospiraceae UCG-001* is associated with greater weight loss in response to lifestyle changes over a 12-month period (**73**). Additionally, a decrease in their abundance could be associated with the development of type 2 diabetes (**74**). While the present study did not evaluate HDL-C levels, research by Lan et al. suggests that reduced butyrate production by *Lachnospiraceae* could contribute to lower HDL-C levels (**75**). Although there is no relevant information regarding *Lachnospiraceae UCG-001* and adiponectin levels, the evidence presented in this study would suggest the importance of this bacterial genus in immunometabolism regulation, making it a target of interest for future research related to microbiota-mediated metabolic modulation. On the other hand, although the *Eubacterium nodatum group*, other SCFA producer, has been associated with periodontitis development, its presence in the intestinal microbiota favors an increase in adiponectin levels mediated or stimulated by an increase in CD45+ CD4+ FOXP3+ IL10+ cells in the VAT. Additionally, it is interesting to note that *Muribaculum* and *Butyricicoccus* exhibited significant modulation of glucose levels through increased IL-10 production by Tregs cells in VAT. An inverse correlation has been demonstrated between metabolites that positively correlate with type 2 diabetes and *Muribaculum* (**76**), while other study showed that the proportion of *Muribaculum intestinale* was significantly decreased in different HFDs (**77**). However, it is paradoxical that *Muribaculum* negatively regulates glucose levels through a decrease in FOXP3+ Tregs abundance in VAT. Although the increase in Tregs cells may be mediated by a direct effect of Omega-3 fatty acids, the available information, cannot explain this finding. On the other hand, there is evidence that *Lachno-clostridium sp.* and *Butyricicoccus sp.* positively correlate with reduced glycemia in diabetic patients treated with glucagon-like peptide-1 receptor agonists (GLP-1 RA) (**78**). Lastly, while a significant correlation has been identified between *Harryflintia* and *Tyzzerella* with Omega-3 fatty acid supplementation in the diet, mediation analyses suggest that these bacterial genera could increase glucose, leptin, and cholesterol levels through the modulation of FOXP3+ Tregs cells in VAT. It is important to note that these observations may be influenced by an overlap of effects. Further studies are required to fully understand the implication of these bacteria in immunometabolic regulation.

In summary, our study has explored the impact of medium-fat diets supplemented with different amount of Omega-3 fatty acids on the interaction between the GM and host tissues, particularly VAT. We demonstrated that constant administration of Omega-3 fatty acids induced specific changes in the composition of the GM, while obesity development was associated with the regulation or induction of Tregs in VAT, with a clear dependence on the dose of Omega-3 administered. Furthermore, our findings suggest a modulatory effect of the microbiota on metabolism, mediated by immunological parameters in VAT. We emphasize the relevance of certain bacterial genera in this effect, which could serve as a starting point for future research. However, it is important to recognize limitations of our study, such as the sample size and the taxonomic analysis that it has been limited to the genus level, which may have hindered the identification of more specific microbial biomarkers for the effect observed.

Despite these limitations, our results lay the groundwork for a better understanding of the complex network of interactions among nutrition, intestinal microbiota, and immunometabolism. Although we have not been able to determine the underlying mechanism of these relationships, our study provides a platform for future research that could explore deeper into this area and open new avenues for the management of obesity and associated metabolic diseases.

## 5. Conclusions

The present results reinforce the importance of Tregs in maintaining local and systemic metabolic balance. Decreased Tregs in VAT have previously been associated with increased inflammation and metabolic dysfunction in obesity models, underscoring the relevance of our observation related to the modulation of Treg levels and IL-10 production in response to Omega-3 supplementation. Additionally, our analyses reveal the possible implication of SCFA in Treg regulation. The complex interaction among the GM, SCFA, and local immune responses highlights the need to delve into the underlying molecular mechanisms. Regarding the microbial profile, we observed an interesting association between certain bacteria and key metabolic parameters. Interestingly, the abundance of *Lachnospiraceae UCG-001* was associated with modulation of leptin, glucose, and cholesterol levels, suggesting a potentially relevant role of this bacterium in metabolic homeostasis. Likewise, we identified the *Eubacterium_nodatum_group* as a possible mediator of adiponectin level regulation, pointing toward the complex interaction between the GM and immunometabolic function.

In summary, our findings support the hypothesis that modulation of the GM composition by Omega-3 fatty acids may be a promising approach in management of obesity and associated metabolic diseases. However, a deeper understanding of the underlying mechanisms and their translation into clinical practice is required before we can fully harness the therapeutic potential of Omega-3 fatty acids.

## Supplementary Materials

Supplementary Table S1A: Supplementary Table S1A: Body weight analysis; Supplementary Table S1B: Delta body weight analysis; Supplementary Table S2: Alpha diversity analysis; Supplementary Table S3: Blood metabolic parameters analysis; Supplementary Table S4: Immune cells population in TAV comparison: Supplementary Table S5A: Correlation between dietary variables and genus with differential abundance among diets; Supplementary Table S5B: Correlation between immune cells population in TAV and genus with differential abundance among diets; Supplementary Table S5C: Correlation between immune cells population in TAV and blood metabolites; Supplementary Table S5D: Correlation between blood metabolites and genus with differential abundance among diets. Supplementary Table S6: Mediation model analysis

### Author Contributions

Conceptualization, N.D.P., R.C.C., and S.A.P.; methodology, N.D.P., N.E., G.B., Y.L.M. and R.C.C.; software, N.D.P. and N.E.; validation, N.D.P., R.C.C., and S.A.P.; formal analysis, N.D.P.; investigation, N.D.P. and S.A.P; resources, C.A.G., R.C.C., M.P.A., and S.A.P.; data curation, N.D.P; writing—original draft preparation, N.D.P. and S.A.P. ; writing—review and editing, C.A.G., R.C.C., M.P.A., and S.A.P.; visualization, N.D.P. and S.A.P.; supervision, R.C.C., M.P.A., and S.A.P.; project administration, R.C.C. and M.P.A.; funding acquisition, C.A.G., R.C.C., M.P.A., and S.A.P.. All authors have read and agreed to the published version of the manuscript.

### Funding

This work was supported in part by funding granted by Secretaría de Investigación y Vinculación Tecnológica from Universidad Católica de Córdoba. Accredited project: Nutritional Quality, a key factor that modulates the composition of the intestinal microbiota and the metabolic immune response in models of obesity. RR 2267-19. This work was supported in part by funding granted by Secretaría de Ciencia y Tecnología from Universidad Nacional de Córdoba (SECyT). Accredited projects: (A) Influence of nutritional quality, innate immunity and acute infection with Trypanosoma cruzi in a diet-induced obesity model. Impact on development/prevention of non-communicable chronic diseases. Amount Res. SECyT-UNC 411/2018. (B) Identification of mononuclear cell metabolic state in T. cruzi infection and its impact on the inflammatory response. SECyT-UNC 113/2018. This work was supported in part by LACE Laboratories SA.

## Institutional Review Board Statement

This research has the authorization of the Institutional Committee for the Care and Use of Laboratory—CICUAL-FCQ—Universidad Nacional de Córdoba, Córdoba, Argentina. RESOLUTION N° 939, EXP-UNC:0023836/2018.

## Informed Consent Statement

Not applicable.

## Data Availability Statement

Sequencing data are accessible in the National Center for Bio-technology Information (NCBI) database under BioProject accession number PRJNA1087685 (https://ncbi.nlm.nih.gov/bioproject/?term=PRJNA1087685)

## Acknowledgments

The authors acknowledge Eduardo Fernandez for continued support. The authors thank Pilar Crespo and Paula Abadie for the technical assistance at the CIBICI-CONICET FACS facility. M.P.A. is a member of the scientific community of Consejo Nacional de Investigaciones Científicas y Técnicas de la República Argentina (CONICET). Y.L.M. and G.B. thank CONICET for the fellowships.

