## Supplementary data for "MICROBIOTA AT THE CROSS ROAD OF DIET AND HEALTH: HOW DIETARY FATS SHAPE BACTERIAL LANDSCAPE AND INFLUENCE GLOBAL HEALTH"

**Supplementary Materials**

**Supplementary Table S1A:** Body weight analysis

| ***Supplementary Table S1A:*** *Body weight analysis* | | | | | | |
| --- | --- | --- | --- | --- | --- | --- |
| **Time** | **Diet** | **Body weight**  **Mean (g)** | **Body weight**  **SD (g)** | **ANOVA p-value** | **Post hoc analysis** | |
|  |  |  |  |  | **Comparison** | **p-value adj** |
| **Week 0** | D1 | 25.4 | 1.4 | 0.552 | - | - |
|  | D2 | 25.4 | 0.6 |  | - | - |
|  | D3 | 26.6 | 2.6 |  | - | - |
| **Time** | **Diet** | **Body weight**  **Mean (g)** | **Body weight**  **SD (g)** | **ANOVA p-value** | **Post hoc analysis** | |
|  |  |  |  |  | **Comparison** | **p-value adj** |
| **Week 4** | D1 | 28.6 | 2.59 | **0.0011** | D1-D2 | **0.0021** |
|  | D2 | 39.3 | 3.1 |  | D1-D3 | **0.0079** |
|  | D3 | 35.5 | 2.4 |  | D2-D3 | 0.1057 |
| **Time** | **Diet** | **Body weight**  **Mean (g)** | **Body weight**  **SD (g)** | **ANOVA p-value** | **Post hoc analysis** | |
|  |  |  |  |  | **Comparison** | **p-value adj** |
| **Week 12** | D1 | 31.8 | 3.3 | **0.0099** | D1-D2 | **0.011** |
|  | D2 | 40.5 | 3.4 |  | D1-D3 | **0.024** |
|  | D3 | 38.3 | 2.8 |  | D2-D3 | 0.373 |
| **Time** | **Diet** | **Body weight**  **Mean (g)** | **Body weight**  **SD (g)** | **ANOVA p-value** | **Post hoc analysis** | |
|  |  |  |  |  | **Comparison** | **p-value adj** |
| **Week 24** | D1 | 37.9 | 3.9 | **0.0166** | D1-D2 | **0.026** |
|  | D2 | 46 | 3.8 |  | D1-D3 | **0.030** |
|  | D3 | 44.9 | 2.1 |  | D2-D3 | 0.641 |
| ***Supplementary Table S1A:*** Body weight analysis for week 0, 4, 12 and 24 in the different diets. Significant differences established using the ANOVA test. Pairwise comparisons using the t-test was utilized in cases of significant results from the ANOVA test (p<0.05). p-values in bold style were statistically significant.  Abbreviations: SD: standard deviation, D1: Diet 1, D2: Diet 2, D3: Diet 3 | | | | | | |

**Supplementary Table S1B:** Delta body weight analysis

| ***Supplementary Table S1B:*** *Delta body weight analysis* | | | | | | |
| --- | --- | --- | --- | --- | --- | --- |
| **Delta** | **Diet** | **Delta body weight**  **Mean (g)** | **Delta body weight**  **SD (g)** | **ANOVA p-value** | **Post hoc analysis** | |
|  |  |  |  |  | **Comparison** | **p-value adj** |
| **Week 4 - Week 0** | D1 | 3.2 | 1.2 | **<0.0001** | D1-D2 | **<0.0001** |
|  | D2 | 13.9 | 2.9 |  | D1-D3 | **0.0073** |
|  | D3 | 8.9 | 1.2 |  | D2-D3 | **0.0163** |
| **Time** | **Diet** | **Delta body weight**  **Mean (g)** | **Delta body weight**  **SD (g)** | **ANOVA p-value** | **Post hoc analysis** | |
|  |  |  |  |  | **Comparison** | **p-value adj** |
| **Week 12 - Week 4** | D1 | 3.2 | 1.8 | 0.136 | D1-D2 | 0.194 |
|  | D2 | 1.2 | 0.9 |  | D1-D3 | 1 |
|  | D3 | 2.9 | 1.1 |  | D2-D3 | 0.362 |
| **Time** | **Diet** | **Delta body weight**  **Mean (g)** | **Delta body weight**  **SD (g)** | **ANOVA p-value** | **Post hoc analysis** | |
|  |  |  |  |  | **Comparison** | **p-value adj** |
| **Week 24 - Week 12** | D1 | 6.1 | 1.4 | 0.799 | D1-D2 | 1 |
|  | D2 | 5.5 | 2 |  | D1-D3 | 1 |
|  | D3 | 6.6 | 3 |  | D2-D3 | 1 |
| ***Supplementary Table S1B:*** Delta body weight analysis for week 4 vs week 0, week 12 vs week 4 and week 24 vs week 12 in the different diets. Significant differences established using the ANOVA test. Pairwise comparisons using the t-test was utilized in cases of significant results from the ANOVA test (p<0.05). p-values in bold style were statistically significant.  Abbreviations: SD: standard deviation, D1: Diet 1, D2: Diet 2, D3: Diet 3 | | | | | | |

**Supplementary Table S2:** Alpha diversity analysis

| ***Supplementary Table S2:*** *Alpha diversity analysis* | | | | | | |
| --- | --- | --- | --- | --- | --- | --- |
| **Alpha diversity index** | **Diet** | **Mean** | **SD** | **Kruskal-Wallis p-value** | **Post hoc analysis** | |
|  |  |  |  |  | **Comparison** | **p-value adj** |
| **Observed ASVs** | D1 | 978 | 41 | 0.055 | - | - |
|  | D2 | 1032 | 17 |  | - | - |
|  | D3 | 930 | 79 |  | - | - |
| **Alpha diversity index** | **Diet** | **Mean** | **SD** | **Kruskal-Wallis p-value** | **Post hoc analysis** | |
|  |  |  |  |  | Comparison | p-value adj |
| **Shannon index** | D1 | 5.48 | 0.08 | **0.018** | D1-D2 | **0.0286** |
|  | D2 | 5.71 | 0.04 |  | D1-D3 | 0.3430 |
|  | D3 | 5.36 | 0.23 |  | D2-D3 | **0.0286** |
| **Alpha diversity index** | **Diet** | **Mean** | **SD** | **Kruskal-Wallis p-value** | **Post hoc analysis** | |
|  |  |  |  |  | **Comparison** | **p-value adj** |
| **Simpson Index** | D1 | 0.989 | 0.001 | **0.037** | D1-D2 | **0.0286** |
|  | D2 | 0.993 | 0.001 |  | D1-D3 | 0.6860 |
|  | D3 | 0.987 | 0.005 |  | D2-D3 | 0.0571 |
| ***Supplementary Table S2:*** Alpha diversity analysis (Observed ASVs, Shannon and Simpson Index) in the different diets. Significant differences established using the Kruskal-Wallis test. Pairwise comparisons using the Wilcoxon signed-rank was utilized in cases of significant results from the Kruskal-Wallis test (p<0.05). p-values in bold style were statistically significant.  Abbreviations: SD: standard deviation, D1: Diet 1, D2: Diet 2, D3: Diet 3 | | | | | | |

**Supplementary Table S3:** Blood metabolic parameters analysis

| **Supplementary Table S3:** Blood metabolic parameters analysis | | | | | | |
| --- | --- | --- | --- | --- | --- | --- |
| **Parameter** | **Diet** | **Mean (mg/dL)** | **SD (mg/dL)** | **ANOVA p-value** |  | |
| **Glucose** | D1 | 138 | 35 | **0.0577** |  |  |
|  | D2 | 239 | 30 |  |  |  |
|  | D3 | 234 | 88 |  |  |  |
| **Parameter** | **Diet** | **Mean (mg/dL)** | **SD (mg/dL)** | **ANOVA p-value** | **Post hoc analysis** | |
|  |  |  |  |  | **Comparison** | **p-value adj** |
| **Cholesterol** | D1 | 78 | 10 | **0.0069** | D1-D2 | **0.0063** |
|  | D2 | 139 | 23 |  | D1-D3 | 0.219 |
|  | D3 | 107 | 25 |  | D2-D3 | 0.156 |
| **Parameter** | **Diet** | **Mean (mg/dL)** | **SD (mg/dL)** | **ANOVA p-value** |  | |
| **Triglycerides** | D1 | 54 | 9 | 0.0544 |  |  |
|  | D2 | 65 | 6 |  |  |  |
|  | D3 | 49 | 8 |  |  |  |
| **Parameter** | **Diet** | **Mean (g/L)** | **SD (g/L)** | **ANOVA p-value** |  | |
| **Total proteins** | D1 | 5.5 | 0.2 | 0.626 |  |  |
|  | D2 | 5.5 | 0.2 |  |  |  |
|  | D3 | 5.3 | 0.5 |  |  |  |
| **Parameter** | **Diet** | **Mean (g/L)** | **SD (g/L)** | **ANOVA p-value** |  |  |
| **Albumin** | D1 | 3.5 | 0.1 | 0.144 |  |  |
|  | D2 | 3.5 | 0.1 |  |  |  |
|  | D3 | 3.3 | 0.1 |  |  |  |
| **Parameter** | **Diet** | **Mean (UI/L)** | **SD (UI/L)** | **Kruskal-Wallis p-value** |  |  |
| **ALT** | D1 | 24 | 2 | 0.101 |  |  |
|  | D2 | 66 | 33 |  |  |  |
|  | D3 | 26 | 12 |  |  |  |
| **Parameter** | **Diet** | **Mean (UI/L)** | **SD (UI/L)** | **ANOVA p-value** |  |  |
| **AST** | D1 | 113 | 45 | 0.787 |  |  |
|  | D2 | 98 | 6 |  |  |  |
|  | D3 | 109 | 29 |  |  |  |
| **Parameter** | **Diet** | **Mean (pg/mL)** | **SD (pg/mL)** | **Kruskal-Wallis p-value** | **Post hoc analysis** | |
|  |  |  |  |  | **Comparison** | **p-value adj** |
| **Leptin** | D1 | 6550 | 1324 | **0.0025** | D1-D2 | **0.0025** |
|  | D2 | 32051 | 5767 |  | D1-D3 | **0.0016** |
|  | D3 | 33917 | 8638 |  | D2-D3 | 0.999 |
| **Parameter** | **Diet** | **Mean (ng/mL)** | **SD (ng/mL)** | **ANOVA p-value** |  |  |
| **Adiponectin** | D1 | 3332 | 964 | 0.206 |  |  |
|  | D2 | 2839 | 1209 |  |  |  |
|  | D3 | 2032 | 588 |  |  |  |
| **Supplementary Table S3:** Blood metabolic parameters comparison for groups D1, D2 and D3 at week 24 of feeding. Significant differences established using ANOVA or Kruskal-Wallis test. Pairwise comparisons using the Wilcoxon signed-rank test were performed if the Krus-kal-Wallis test yielded a significant result, while the t-test was utilized in cases of signifi-cant results from the ANOVA test (p<0.05). p-values in bold style were statistically significant.  Abbreviations: SD: standard deviation, D1: Diet 1, D2: Diet 2, D3: Diet 3, AST: aspartate aminotransferase, ALT: alanine aminotransferase. | | | | | | |

**Supplementary Table S4:** Immune cells population in TAV comparison

| ***Supplementary Table S4:*** *Immune cells population in TAV comparison* | | | | | | |
| --- | --- | --- | --- | --- | --- | --- |
| **Parameter** | Diet | Mean (%) | SD (%) | ANOVA p-value |  | |
| **% CD45+ CD4+** | D1 | 12.4 | 2.6 | 0.604 |  |  |
|  | D2 | 14 | 3.9 |  |  |  |
|  | D3 | 16 | 7 |  |  |  |
| **Parameter** | **Diet** | **Mean (%)** | **SD (%)** | **ANOVA p-value** | **Post hoc analysis** | |
|  |  |  |  |  | **Comparison** | **p-value adj** |
| **% CD45+ CD4+ FOXP3+** | D1 | 7.8 | 1.9 | **0.0002** | D1-D2 | **0.0002** |
|  | D2 | 26.8 | 5.5 |  | D1-D3 | **0.0051** |
|  | D3 | 19.4 | 2.8 |  | D2-D3 | 0.0624 |
| **Parameter** | **Diet** | **Mean (%)** | **SD (%)** | **ANOVA p-value** | **Post hoc analysis** | |
|  |  |  |  |  | **Comparison** | **p-value adj** |
| **% CD45+ CD4+ FOXP3+ IL-10+** | D1 | 79.9 | 13.1 | **0.0224** | D1-D2 | **0.0341** |
|  | D2 | 55.3 | 2.7 |  | D1-D3 | 0.0988 |
|  | D3 | 60.4 | 13.6 |  | D2-D3 | 1 |
| **Parameter** | **Diet** | **Mean (AU)** | **SD (AU)** | **Kruskal-Wallis p-value** | **Post hoc analysis** | |
|  |  |  |  |  | **Comparison** | **p-value adj** |
| **MFI (CD45+ CD4+ FOXP3+ IL-10+)** | D1 | 331 | 201 | **0.0210** | D1-D2 | **0.029** |
|  | D2 | 78 | 8 |  | D1-D3 | **0.029** |
|  | D3 | 90 | 22 |  | D2-D3 | 0.486 |
| **Parameter** | **Diet** | **Mean (%)** | **SD (%)** | **ANOVA p-value** |  |  |
| **% CD45+ CD11b+** | D1 | 20.7 | 7.2 | 0.782 |  |  |
|  | D2 | 21.7 | 4.4 |  |  |  |
|  | D3 | 24.1 | 8.6 |  |  |  |
| **Parameter** | **Diet** | **Mean (%)** | **SD (%)** | **ANOVA p-value** |  |  |
| **% CD45+ CD11b+ F4/80+** | D1 | 26 | 7.9 | 0.632 |  |  |
|  | D2 | 21.3 | 3.2 |  |  |  |
|  | D3 | 24.4 | 8.5 |  |  |  |
| **Supplementary Table S4:** Immune cells population in TAV comparison for groups D1, D2 and D3 at week 24 of feeding. Significant differences established using ANOVA or Kruskal-Wallis test. Pairwise comparisons using the Wilcoxon signed-rank test were performed if the Krus-kal-Wallis test yielded a significant result, while the t-test was utilized in cases of signifi-cant results from the ANOVA test (p<0.05). p-values in bold style were statistically significant.  Abbreviations: SD: standard deviation, D1: Diet 1, D2: Diet 2, D3: Diet 3, CD: cluster of differentiation, MFI: median fluorescence intensity. | | | | | | |

**Supplementary Table S5A:** Correlation between dietary variables and genus with differential abundance among diets

| ***Supplementary Table S5A*** | | | |
| --- | --- | --- | --- |
| **Dietary variable** | **Genus** | **rho** | **p-value** |
| Total Fat content | *Alloprevotella* | -0.65 | 0.0209 |
| Total Fat content | *Prevotellaceae UCG-001* | -0.37 | 0.2405 |
| Total Fat content | *Lachnoanaerobaculum* | 0.68 | 0.0159 |
| Total Fat content | *Helicobacter* | 0.31 | 0.3243 |
| Total Fat content | *Candidatus Saccharimonas* | -0.43 | 0.1605 |
| Total Fat content | *Flavonifractor* | -0.20 | 0.5436 |
| Total Fat content | *Parasutterella* | -0.60 | 0.0373 |
| Total Fat content | *Odoribacter* | 0.67 | 0.0164 |
| Total Fat content | *Mucispirillum* | 0.40 | 0.1931 |
| Total Fat content | *Oscillospira* | 0.31 | 0.3249 |
| Total Fat content | *Muribaculum* | -0.86 | 0.0003 |
| Total Fat content | *Herbinix* | -0.04 | 0.8971 |
| Total Fat content | *Negativibacillus* | 0.58 | 0.0479 |
| Total Fat content | *[Ruminococcus] gnavus group* | 0.62 | 0.0327 |
| Total Fat content | *Oscillibacter* | 0.29 | 0.3686 |
| Total Fat content | *Anaerotruncus* | 0.59 | 0.0426 |
| Total Fat content | *Alistipes* | 0.45 | 0.1402 |
| Total Fat content | *Prevotella_7* | -0.70 | 0.0112 |
| Total Fat content | *Harryflintia* | 0.12 | 0.7005 |
| Total Fat content | *Anaeroplasma* | -0.82 | 0.0011 |
| Total Fat content | *Parabacteroides* | -0.47 | 0.1242 |
| Total Fat content | *UCG-001* | -0.63 | 0.0294 |
| Total Fat content | *Butyricicoccus* | -0.84 | 0.0007 |
| Total Fat content | *Stomatobaculum* | 0.11 | 0.7316 |
| Total Fat content | *Prevotellaceae NK3B31 group* | -0.03 | 0.9385 |
| Total Fat content | *Lactobacillus* | -0.65 | 0.0224 |
| Total Fat content | *[Eubacterium] eligens group* | 0.59 | 0.0436 |
| Total Fat content | *Angelakisella* | 0.52 | 0.0853 |
| Total Fat content | *ASF356* | -0.18 | 0.5841 |
| Total Fat content | *[Eubacterium] nodatum group* | -0.74 | 0.0064 |
| Total Fat content | *Roseburia* | 0.39 | 0.2137 |
| Total Fat content | *Erysipelatoclostridium* | -0.28 | 0.3862 |
| Total Fat content | *Paludicola* | -0.53 | 0.0734 |
| Total Fat content | *Rikenella* | 0.47 | 0.1214 |
| Total Fat content | *Lachnospiraceae UCG-001* | -0.07 | 0.8218 |
| Total Fat content | *Tyzzerella* | 0.48 | 0.1151 |
| Absolute Omega-3 content | *Alloprevotella* | -0.69 | 0.0129 |
| Absolute Omega-3 content | *Prevotellaceae UCG-001* | -0.58 | 0.0501 |
| Absolute Omega-3 content | *Lachnoanaerobaculum* | 0.54 | 0.0705 |
| Absolute Omega-3 content | *Helicobacter* | -0.04 | 0.9083 |
| Absolute Omega-3 content | *Candidatus Saccharimonas* | -0.14 | 0.6587 |
| Absolute Omega-3 content | *Flavonifractor* | 0.24 | 0.4458 |
| Absolute Omega-3 content | *Parasutterella* | -0.53 | 0.0742 |
| Absolute Omega-3 content | *Odoribacter* | 0.74 | 0.0061 |
| Absolute Omega-3 content | *Mucispirillum* | 0.31 | 0.3203 |
| Absolute Omega-3 content | *Oscillospira* | 0.63 | 0.0270 |
| Absolute Omega-3 content | *Muribaculum* | -0.86 | 0.0003 |
| Absolute Omega-3 content | *Herbinix* | -0.24 | 0.4488 |
| Absolute Omega-3 content | *Negativibacillus* | 0.42 | 0.1772 |
| Absolute Omega-3 content | *[Ruminococcus] gnavus group* | 0.85 | 0.0004 |
| Absolute Omega-3 content | *Oscillibacter* | 0.74 | 0.0058 |
| Absolute Omega-3 content | *Anaerotruncus* | 0.52 | 0.0852 |
| Absolute Omega-3 content | *Alistipes* | 0.69 | 0.0134 |
| Absolute Omega-3 content | *Prevotella_7* | -0.62 | 0.0333 |
| Absolute Omega-3 content | *Harryflintia* | 0.53 | 0.0745 |
| Absolute Omega-3 content | *Anaeroplasma* | -0.72 | 0.0086 |
| Absolute Omega-3 content | *Parabacteroides* | 0.09 | 0.7698 |
| Absolute Omega-3 content | *UCG-001* | -0.53 | 0.0764 |
| Absolute Omega-3 content | *Butyricicoccus* | -0.60 | 0.0396 |
| Absolute Omega-3 content | *Stomatobaculum* | 0.48 | 0.1119 |
| Absolute Omega-3 content | *Prevotellaceae NK3B31 group* | -0.34 | 0.2743 |
| Absolute Omega-3 content | *Lactobacillus* | -0.48 | 0.1105 |
| Absolute Omega-3 content | *[Eubacterium] eligens group* | 0.45 | 0.1407 |
| Absolute Omega-3 content | *Angelakisella* | 0.17 | 0.6006 |
| Absolute Omega-3 content | *ASF356* | 0.44 | 0.1538 |
| Absolute Omega-3 content | *[Eubacterium] nodatum group* | -0.62 | 0.0298 |
| Absolute Omega-3 content | *Roseburia* | 0.26 | 0.4106 |
| Absolute Omega-3 content | *Erysipelatoclostridium* | -0.52 | 0.0808 |
| Absolute Omega-3 content | *Paludicola* | -0.13 | 0.6802 |
| Absolute Omega-3 content | *Rikenella* | 0.68 | 0.0152 |
| Absolute Omega-3 content | *Lachnospiraceae UCG-001* | -0.45 | 0.1429 |
| Absolute Omega-3 content | *Tyzzerella* | 0.91 | 0.0000 |
| Absolute Omega-6 content | *Alloprevotella* | -0.59 | 0.0454 |
| Absolute Omega-6 content | *Prevotellaceae UCG-001* | -0.27 | 0.3874 |
| Absolute Omega-6 content | *Lachnoanaerobaculum* | 0.65 | 0.0212 |
| Absolute Omega-6 content | *Helicobacter* | 0.38 | 0.2198 |
| Absolute Omega-6 content | *Candidatus Saccharimonas* | -0.48 | 0.1181 |
| Absolute Omega-6 content | *Flavonifractor* | -0.30 | 0.3403 |
| Absolute Omega-6 content | *Parasutterella* | -0.57 | 0.0528 |
| Absolute Omega-6 content | *Odoribacter* | 0.59 | 0.0417 |
| Absolute Omega-6 content | *Mucispirillum* | 0.39 | 0.2065 |
| Absolute Omega-6 content | *Oscillospira* | 0.19 | 0.5508 |
| Absolute Omega-6 content | *Muribaculum* | -0.79 | 0.0025 |
| Absolute Omega-6 content | *Herbinix* | 0.02 | 0.9542 |
| Absolute Omega-6 content | *Negativibacillus* | 0.57 | 0.0508 |
| Absolute Omega-6 content | *[Ruminococcus] gnavus group* | 0.49 | 0.1023 |
| Absolute Omega-6 content | *Oscillibacter* | 0.13 | 0.6866 |
| Absolute Omega-6 content | *Anaerotruncus* | 0.56 | 0.0583 |
| Absolute Omega-6 content | *Alistipes* | 0.34 | 0.2731 |
| Absolute Omega-6 content | *Prevotella_7* | -0.66 | 0.0192 |
| Absolute Omega-6 content | *Harryflintia* | 0.00 | 0.9927 |
| Absolute Omega-6 content | *Anaeroplasma* | -0.78 | 0.0030 |
| Absolute Omega-6 content | *Parabacteroides* | -0.59 | 0.0450 |
| Absolute Omega-6 content | *UCG-001* | -0.60 | 0.0405 |
| Absolute Omega-6 content | *Butyricicoccus* | -0.83 | 0.0009 |
| Absolute Omega-6 content | *Stomatobaculum* | 0.00 | 0.9884 |
| Absolute Omega-6 content | *Prevotellaceae NK3B31 group* | 0.07 | 0.8345 |
| Absolute Omega-6 content | *Lactobacillus* | -0.64 | 0.0258 |
| Absolute Omega-6 content | *[Eubacterium] eligens group* | 0.58 | 0.0501 |
| Absolute Omega-6 content | *Angelakisella* | 0.57 | 0.0534 |
| Absolute Omega-6 content | *ASF356* | -0.33 | 0.2876 |
| Absolute Omega-6 content | *[Eubacterium] nodatum group* | -0.70 | 0.0111 |
| Absolute Omega-6 content | *Roseburia* | 0.39 | 0.2129 |
| Absolute Omega-6 content | *Erysipelatoclostridium* | -0.18 | 0.5751 |
| Absolute Omega-6 content | *Paludicola* | -0.60 | 0.0391 |
| Absolute Omega-6 content | *Rikenella* | 0.37 | 0.2356 |
| Absolute Omega-6 content | *Lachnospiraceae UCG-001* | 0.04 | 0.9007 |
| Absolute Omega-6 content | *Tyzzerella* | 0.31 | 0.3209 |
| Omega-3/Omega-6 | *Alloprevotella* | -0.61 | 0.0366 |
| Omega-3/Omega-6 | *Prevotellaceae UCG-001* | -0.58 | 0.0484 |
| Omega-3/Omega-6 | *Lachnoanaerobaculum* | 0.41 | 0.1910 |
| Omega-3/Omega-6 | *Helicobacter* | -0.18 | 0.5844 |
| Omega-3/Omega-6 | *Candidatus Saccharimonas* | 0.00 | 0.9940 |
| Omega-3/Omega-6 | *Flavonifractor* | 0.39 | 0.2107 |
| Omega-3/Omega-6 | *Parasutterella* | -0.43 | 0.1658 |
| Omega-3/Omega-6 | *Odoribacter* | 0.66 | 0.0197 |
| Omega-3/Omega-6 | *Mucispirillum* | 0.23 | 0.4682 |
| Omega-3/Omega-6 | *Oscillospira* | 0.68 | 0.0158 |
| Omega-3/Omega-6 | *Muribaculum* | -0.74 | 0.0063 |
| Omega-3/Omega-6 | *Herbinix* | -0.29 | 0.3610 |
| Omega-3/Omega-6 | *Negativibacillus* | 0.29 | 0.3603 |
| Omega-3/Omega-6 | *[Ruminococcus] gnavus group* | 0.83 | 0.0009 |
| Omega-3/Omega-6 | *Oscillibacter* | 0.82 | 0.0010 |
| Omega-3/Omega-6 | *Anaerotruncus* | 0.41 | 0.1834 |
| Omega-3/Omega-6 | *Alistipes* | 0.69 | 0.0136 |
| Omega-3/Omega-6 | *Prevotella_7* | -0.49 | 0.1044 |
| Omega-3/Omega-6 | *Harryflintia* | 0.63 | 0.0297 |
| Omega-3/Omega-6 | *Anaeroplasma* | -0.57 | 0.0517 |
| Omega-3/Omega-6 | *Parabacteroides* | 0.31 | 0.3207 |
| Omega-3/Omega-6 | *UCG-001* | -0.41 | 0.1805 |
| Omega-3/Omega-6 | *Butyricicoccus* | -0.42 | 0.1794 |
| Omega-3/Omega-6 | *Stomatobaculum* | 0.57 | 0.0545 |
| Omega-3/Omega-6 | *Prevotellaceae NK3B31 group* | -0.43 | 0.1676 |
| Omega-3/Omega-6 | *Lactobacillus* | -0.35 | 0.2692 |
| Omega-3/Omega-6 | *[Eubacterium] eligens group* | 0.33 | 0.2953 |
| Omega-3/Omega-6 | *Angelakisella* | 0.00 | 0.9986 |
| Omega-3/Omega-6 | *ASF356* | 0.63 | 0.0283 |
| Omega-3/Omega-6 | *[Eubacterium] nodatum group* | -0.49 | 0.1062 |
| Omega-3/Omega-6 | *Roseburia* | 0.17 | 0.5911 |
| Omega-3/Omega-6 | *Erysipelatoclostridium* | -0.55 | 0.0636 |
| Omega-3/Omega-6 | *Paludicola* | 0.05 | 0.8732 |
| Omega-3/Omega-6 | *Rikenella* | 0.67 | 0.0178 |
| Omega-3/Omega-6 | *Lachnospiraceae UCG-001* | -0.54 | 0.0698 |
| Omega-3/Omega-6 | *Tyzzerella* | 0.96 | <0.0001 |
| **Supplementary Table S5A:** Correlation between dietary variables and Genus with differential abundance among diets. Correlation significant established using the Pearson test (Significant p-value<0.05 was consider). | | | |

**Supplementary Table S5B:** Correlation between immune cells population in TAV and genus with differential abundance among diets

| **Supplementary Table S5B** | | | |
| --- | --- | --- | --- |
| **Genus** | **Immune cells variable in TAV** | **rho** | **p-value** |
| Alloprevotella | % CD45+ CD4+ | 0.08 | 0.8082 |
| Prevotellaceae UCG-001 | % CD45+ CD4+ | 0.57 | 0.0544 |
| Lachnoanaerobaculum | % CD45+ CD4+ | 0.05 | 0.8658 |
| Helicobacter | % CD45+ CD4+ | 0.09 | 0.7854 |
| Candidatus Saccharimonas | % CD45+ CD4+ | -0.37 | 0.2370 |
| Flavonifractor | % CD45+ CD4+ | -0.20 | 0.5389 |
| Parasutterella | % CD45+ CD4+ | -0.05 | 0.8788 |
| Odoribacter | % CD45+ CD4+ | -0.23 | 0.4766 |
| Mucispirillum | % CD45+ CD4+ | -0.20 | 0.5290 |
| Oscillospira | % CD45+ CD4+ | 0.29 | 0.3622 |
| Muribaculum | % CD45+ CD4+ | -0.27 | 0.3906 |
| Herbinix | % CD45+ CD4+ | -0.18 | 0.5773 |
| Negativibacillus | % CD45+ CD4+ | -0.23 | 0.4797 |
| [Ruminococcus] gnavus group | % CD45+ CD4+ | -0.01 | 0.9636 |
| Oscillibacter | % CD45+ CD4+ | 0.13 | 0.6852 |
| Anaerotruncus | % CD45+ CD4+ | 0.18 | 0.5808 |
| Alistipes | % CD45+ CD4+ | -0.33 | 0.2914 |
| Prevotella_7 | % CD45+ CD4+ | 0.28 | 0.3823 |
| Harryflintia | % CD45+ CD4+ | 0.01 | 0.9862 |
| Anaeroplasma | % CD45+ CD4+ | -0.14 | 0.6688 |
| Parabacteroides | % CD45+ CD4+ | -0.03 | 0.9141 |
| UCG-001 | % CD45+ CD4+ | -0.16 | 0.6154 |
| Butyricicoccus | % CD45+ CD4+ | -0.38 | 0.2238 |
| Stomatobaculum | % CD45+ CD4+ | -0.26 | 0.4124 |
| Prevotellaceae NK3B31 group | % CD45+ CD4+ | 0.73 | 0.0074 |
| Lactobacillus | % CD45+ CD4+ | -0.28 | 0.3702 |
| [Eubacterium] eligens group | % CD45+ CD4+ | 0.25 | 0.4411 |
| Angelakisella | % CD45+ CD4+ | -0.23 | 0.4767 |
| ASF356 | % CD45+ CD4+ | -0.01 | 0.9680 |
| [Eubacterium] nodatum group | % CD45+ CD4+ | -0.15 | 0.6390 |
| Roseburia | % CD45+ CD4+ | 0.03 | 0.9357 |
| Erysipelatoclostridium | % CD45+ CD4+ | 0.20 | 0.5262 |
| Paludicola | % CD45+ CD4+ | -0.25 | 0.4414 |
| Rikenella | % CD45+ CD4+ | -0.14 | 0.6593 |
| Lachnospiraceae UCG-001 | % CD45+ CD4+ | -0.08 | 0.7994 |
| Tyzzerella | % CD45+ CD4+ | -0.13 | 0.6804 |
| Alloprevotella | % CD45+ CD4+ FOXP3+ | -0.66 | 0.0183 |
| Prevotellaceae UCG-001 | % CD45+ CD4+ FOXP3+ | -0.60 | 0.0381 |
| Lachnoanaerobaculum | % CD45+ CD4+ FOXP3+ | 0.54 | 0.0680 |
| Helicobacter | % CD45+ CD4+ FOXP3+ | 0.12 | 0.7188 |
| Candidatus Saccharimonas | % CD45+ CD4+ FOXP3+ | -0.08 | 0.8116 |
| Flavonifractor | % CD45+ CD4+ FOXP3+ | -0.01 | 0.9777 |
| Parasutterella | % CD45+ CD4+ FOXP3+ | -0.58 | 0.0485 |
| Odoribacter | % CD45+ CD4+ FOXP3+ | 0.76 | 0.0038 |
| Mucispirillum | % CD45+ CD4+ FOXP3+ | 0.38 | 0.2223 |
| Oscillospira | % CD45+ CD4+ FOXP3+ | 0.40 | 0.1960 |
| Muribaculum | % CD45+ CD4+ FOXP3+ | -0.87 | 0.0003 |
| Herbinix | % CD45+ CD4+ FOXP3+ | -0.19 | 0.5448 |
| Negativibacillus | % CD45+ CD4+ FOXP3+ | 0.58 | 0.0489 |
| [Ruminococcus] gnavus group | % CD45+ CD4+ FOXP3+ | 0.78 | 0.0026 |
| Oscillibacter | % CD45+ CD4+ FOXP3+ | 0.45 | 0.1400 |
| Anaerotruncus | % CD45+ CD4+ FOXP3+ | 0.49 | 0.1065 |
| Alistipes | % CD45+ CD4+ FOXP3+ | 0.61 | 0.0355 |
| Prevotella_7 | % CD45+ CD4+ FOXP3+ | -0.76 | 0.0045 |
| Harryflintia | % CD45+ CD4+ FOXP3+ | 0.43 | 0.1639 |
| Anaeroplasma | % CD45+ CD4+ FOXP3+ | -0.73 | 0.0068 |
| Parabacteroides | % CD45+ CD4+ FOXP3+ | -0.17 | 0.5939 |
| UCG-001 | % CD45+ CD4+ FOXP3+ | -0.59 | 0.0452 |
| Butyricicoccus | % CD45+ CD4+ FOXP3+ | -0.61 | 0.0344 |
| Stomatobaculum | % CD45+ CD4+ FOXP3+ | 0.37 | 0.2361 |
| Prevotellaceae NK3B31 group | % CD45+ CD4+ FOXP3+ | -0.28 | 0.3838 |
| Lactobacillus | % CD45+ CD4+ FOXP3+ | -0.59 | 0.0444 |
| [Eubacterium] eligens group | % CD45+ CD4+ FOXP3+ | 0.38 | 0.2285 |
| Angelakisella | % CD45+ CD4+ FOXP3+ | 0.39 | 0.2093 |
| ASF356 | % CD45+ CD4+ FOXP3+ | 0.12 | 0.7218 |
| [Eubacterium] nodatum group | % CD45+ CD4+ FOXP3+ | -0.67 | 0.0169 |
| Roseburia | % CD45+ CD4+ FOXP3+ | 0.40 | 0.2011 |
| Erysipelatoclostridium | % CD45+ CD4+ FOXP3+ | -0.47 | 0.1265 |
| Paludicola | % CD45+ CD4+ FOXP3+ | -0.37 | 0.2325 |
| Rikenella | % CD45+ CD4+ FOXP3+ | 0.78 | 0.0030 |
| Lachnospiraceae UCG-001 | % CD45+ CD4+ FOXP3+ | -0.22 | 0.5012 |
| Tyzzerella | % CD45+ CD4+ FOXP3+ | 0.79 | 0.0024 |
| Alloprevotella | % CD45+ CD4+ FOXP3+ IL-10+ | 0.39 | 0.2041 |
| Prevotellaceae UCG-001 | % CD45+ CD4+ FOXP3+ IL-10+ | 0.47 | 0.1228 |
| Lachnoanaerobaculum | % CD45+ CD4+ FOXP3+ IL-10+ | -0.20 | 0.5435 |
| Helicobacter | % CD45+ CD4+ FOXP3+ IL-10+ | -0.37 | 0.2325 |
| Candidatus Saccharimonas | % CD45+ CD4+ FOXP3+ IL-10+ | 0.23 | 0.4696 |
| Flavonifractor | % CD45+ CD4+ FOXP3+ IL-10+ | -0.13 | 0.6868 |
| Parasutterella | % CD45+ CD4+ FOXP3+ IL-10+ | 0.39 | 0.2155 |
| Odoribacter | % CD45+ CD4+ FOXP3+ IL-10+ | -0.46 | 0.1332 |
| Mucispirillum | % CD45+ CD4+ FOXP3+ IL-10+ | -0.50 | 0.0963 |
| Oscillospira | % CD45+ CD4+ FOXP3+ IL-10+ | -0.51 | 0.0901 |
| Muribaculum | % CD45+ CD4+ FOXP3+ IL-10+ | 0.52 | 0.0817 |
| Herbinix | % CD45+ CD4+ FOXP3+ IL-10+ | 0.55 | 0.0651 |
| Negativibacillus | % CD45+ CD4+ FOXP3+ IL-10+ | -0.06 | 0.8455 |
| [Ruminococcus] gnavus group | % CD45+ CD4+ FOXP3+ IL-10+ | -0.54 | 0.0711 |
| Oscillibacter | % CD45+ CD4+ FOXP3+ IL-10+ | -0.37 | 0.2337 |
| Anaerotruncus | % CD45+ CD4+ FOXP3+ IL-10+ | -0.66 | 0.0196 |
| Alistipes | % CD45+ CD4+ FOXP3+ IL-10+ | -0.39 | 0.2146 |
| Prevotella_7 | % CD45+ CD4+ FOXP3+ IL-10+ | 0.72 | 0.0086 |
| Harryflintia | % CD45+ CD4+ FOXP3+ IL-10+ | -0.10 | 0.7550 |
| Anaeroplasma | % CD45+ CD4+ FOXP3+ IL-10+ | 0.64 | 0.0252 |
| Parabacteroides | % CD45+ CD4+ FOXP3+ IL-10+ | 0.47 | 0.1222 |
| UCG-001 | % CD45+ CD4+ FOXP3+ IL-10+ | 0.37 | 0.2357 |
| Butyricicoccus | % CD45+ CD4+ FOXP3+ IL-10+ | 0.63 | 0.0286 |
| Stomatobaculum | % CD45+ CD4+ FOXP3+ IL-10+ | -0.12 | 0.7081 |
| Prevotellaceae NK3B31 group | % CD45+ CD4+ FOXP3+ IL-10+ | -0.05 | 0.8827 |
| Lactobacillus | % CD45+ CD4+ FOXP3+ IL-10+ | 0.25 | 0.4372 |
| [Eubacterium] eligens group | % CD45+ CD4+ FOXP3+ IL-10+ | -0.61 | 0.0344 |
| Angelakisella | % CD45+ CD4+ FOXP3+ IL-10+ | -0.14 | 0.6556 |
| ASF356 | % CD45+ CD4+ FOXP3+ IL-10+ | 0.08 | 0.7959 |
| [Eubacterium] nodatum group | % CD45+ CD4+ FOXP3+ IL-10+ | 0.60 | 0.0406 |
| Roseburia | % CD45+ CD4+ FOXP3+ IL-10+ | -0.57 | 0.0511 |
| Erysipelatoclostridium | % CD45+ CD4+ FOXP3+ IL-10+ | 0.41 | 0.1834 |
| Paludicola | % CD45+ CD4+ FOXP3+ IL-10+ | 0.12 | 0.7154 |
| Rikenella | % CD45+ CD4+ FOXP3+ IL-10+ | -0.46 | 0.1288 |
| Lachnospiraceae UCG-001 | % CD45+ CD4+ FOXP3+ IL-10+ | 0.60 | 0.0378 |
| Tyzzerella | % CD45+ CD4+ FOXP3+ IL-10+ | -0.47 | 0.1217 |
| Alloprevotella | MFI (CD45+ CD4+ FOXP3+ IL-10+) | 0.30 | 0.3402 |
| Prevotellaceae UCG-001 | MFI (CD45+ CD4+ FOXP3+ IL-10+) | 0.32 | 0.3103 |
| Lachnoanaerobaculum | MFI (CD45+ CD4+ FOXP3+ IL-10+) | -0.46 | 0.1311 |
| Helicobacter | MFI (CD45+ CD4+ FOXP3+ IL-10+) | -0.27 | 0.4049 |
| Candidatus Saccharimonas | MFI (CD45+ CD4+ FOXP3+ IL-10+) | 0.14 | 0.6682 |
| Flavonifractor | MFI (CD45+ CD4+ FOXP3+ IL-10+) | 0.02 | 0.9429 |
| Parasutterella | MFI (CD45+ CD4+ FOXP3+ IL-10+) | 0.45 | 0.1382 |
| Odoribacter | MFI (CD45+ CD4+ FOXP3+ IL-10+) | -0.52 | 0.0837 |
| Mucispirillum | MFI (CD45+ CD4+ FOXP3+ IL-10+) | -0.41 | 0.1887 |
| Oscillospira | MFI (CD45+ CD4+ FOXP3+ IL-10+) | -0.42 | 0.1741 |
| Muribaculum | MFI (CD45+ CD4+ FOXP3+ IL-10+) | 0.66 | 0.0196 |
| Herbinix | MFI (CD45+ CD4+ FOXP3+ IL-10+) | 0.13 | 0.6880 |
| Negativibacillus | MFI (CD45+ CD4+ FOXP3+ IL-10+) | -0.40 | 0.1982 |
| [Ruminococcus] gnavus group | MFI (CD45+ CD4+ FOXP3+ IL-10+) | -0.54 | 0.0679 |
| Oscillibacter | MFI (CD45+ CD4+ FOXP3+ IL-10+) | -0.37 | 0.2330 |
| Anaerotruncus | MFI (CD45+ CD4+ FOXP3+ IL-10+) | -0.48 | 0.1143 |
| Alistipes | MFI (CD45+ CD4+ FOXP3+ IL-10+) | -0.42 | 0.1744 |
| Prevotella_7 | MFI (CD45+ CD4+ FOXP3+ IL-10+) | 0.69 | 0.0122 |
| Harryflintia | MFI (CD45+ CD4+ FOXP3+ IL-10+) | 0.00 | 0.9973 |
| Anaeroplasma | MFI (CD45+ CD4+ FOXP3+ IL-10+) | 0.80 | 0.0019 |
| Parabacteroides | MFI (CD45+ CD4+ FOXP3+ IL-10+) | 0.53 | 0.0762 |
| UCG-001 | MFI (CD45+ CD4+ FOXP3+ IL-10+) | 0.33 | 0.2985 |
| Butyricicoccus | MFI (CD45+ CD4+ FOXP3+ IL-10+) | 0.65 | 0.0223 |
| Stomatobaculum | MFI (CD45+ CD4+ FOXP3+ IL-10+) | 0.25 | 0.4349 |
| Prevotellaceae NK3B31 group | MFI (CD45+ CD4+ FOXP3+ IL-10+) | -0.04 | 0.8952 |
| Lactobacillus | MFI (CD45+ CD4+ FOXP3+ IL-10+) | 0.37 | 0.2431 |
| [Eubacterium] eligens group | MFI (CD45+ CD4+ FOXP3+ IL-10+) | -0.46 | 0.1318 |
| Angelakisella | MFI (CD45+ CD4+ FOXP3+ IL-10+) | -0.38 | 0.2257 |
| ASF356 | MFI (CD45+ CD4+ FOXP3+ IL-10+) | 0.10 | 0.7673 |
| [Eubacterium] nodatum group | MFI (CD45+ CD4+ FOXP3+ IL-10+) | 0.62 | 0.0325 |
| Roseburia | MFI (CD45+ CD4+ FOXP3+ IL-10+) | -0.31 | 0.3241 |
| Erysipelatoclostridium | MFI (CD45+ CD4+ FOXP3+ IL-10+) | 0.37 | 0.2375 |
| Paludicola | MFI (CD45+ CD4+ FOXP3+ IL-10+) | 0.24 | 0.4555 |
| Rikenella | MFI (CD45+ CD4+ FOXP3+ IL-10+) | -0.27 | 0.3984 |
| Lachnospiraceae UCG-001 | MFI (CD45+ CD4+ FOXP3+ IL-10+) | 0.14 | 0.6625 |
| Tyzzerella | MFI (CD45+ CD4+ FOXP3+ IL-10+) | -0.39 | 0.2049 |
| Alloprevotella | % CD45+ CD11b+ | -0.35 | 0.2709 |
| Prevotellaceae UCG-001 | % CD45+ CD11b+ | 0.04 | 0.9036 |
| Lachnoanaerobaculum | % CD45+ CD11b+ | 0.14 | 0.6729 |
| Helicobacter | % CD45+ CD11b+ | 0.61 | 0.0357 |
| Candidatus Saccharimonas | % CD45+ CD11b+ | -0.49 | 0.1041 |
| Flavonifractor | % CD45+ CD11b+ | -0.18 | 0.5747 |
| Parasutterella | % CD45+ CD11b+ | -0.16 | 0.6197 |
| Odoribacter | % CD45+ CD11b+ | -0.17 | 0.6003 |
| Mucispirillum | % CD45+ CD11b+ | 0.40 | 0.1997 |
| Oscillospira | % CD45+ CD11b+ | -0.18 | 0.5660 |
| Muribaculum | % CD45+ CD11b+ | 0.16 | 0.6115 |
| Herbinix | % CD45+ CD11b+ | 0.09 | 0.7884 |
| Negativibacillus | % CD45+ CD11b+ | 0.07 | 0.8383 |
| [Ruminococcus] gnavus group | % CD45+ CD11b+ | 0.05 | 0.8701 |
| Oscillibacter | % CD45+ CD11b+ | -0.19 | 0.5566 |
| Anaerotruncus | % CD45+ CD11b+ | 0.53 | 0.0770 |
| Alistipes | % CD45+ CD11b+ | -0.47 | 0.1237 |
| Prevotella_7 | % CD45+ CD11b+ | -0.16 | 0.6116 |
| Harryflintia | % CD45+ CD11b+ | 0.04 | 0.9069 |
| Anaeroplasma | % CD45+ CD11b+ | -0.17 | 0.5949 |
| Parabacteroides | % CD45+ CD11b+ | -0.31 | 0.3271 |
| UCG-001 | % CD45+ CD11b+ | -0.15 | 0.6493 |
| Butyricicoccus | % CD45+ CD11b+ | 0.09 | 0.7875 |
| Stomatobaculum | % CD45+ CD11b+ | 0.35 | 0.2608 |
| Prevotellaceae NK3B31 group | % CD45+ CD11b+ | -0.11 | 0.7436 |
| Lactobacillus | % CD45+ CD11b+ | -0.14 | 0.6533 |
| [Eubacterium] eligens group | % CD45+ CD11b+ | 0.59 | 0.0452 |
| Angelakisella | % CD45+ CD11b+ | 0.42 | 0.1759 |
| ASF356 | % CD45+ CD11b+ | -0.32 | 0.3098 |
| [Eubacterium] nodatum group | % CD45+ CD11b+ | -0.06 | 0.8418 |
| Roseburia | % CD45+ CD11b+ | 0.62 | 0.0324 |
| Erysipelatoclostridium | % CD45+ CD11b+ | 0.02 | 0.9590 |
| Paludicola | % CD45+ CD11b+ | -0.21 | 0.5101 |
| Rikenella | % CD45+ CD11b+ | -0.17 | 0.5995 |
| Lachnospiraceae UCG-001 | % CD45+ CD11b+ | -0.22 | 0.4908 |
| Tyzzerella | % CD45+ CD11b+ | -0.08 | 0.8041 |
| Alloprevotella | % CD45+ CD11b+ F4/80+ | 0.41 | 0.1871 |
| Prevotellaceae UCG-001 | % CD45+ CD11b+ F4/80+ | 0.44 | 0.1543 |
| Lachnoanaerobaculum | % CD45+ CD11b+ F4/80+ | 0.25 | 0.4383 |
| Helicobacter | % CD45+ CD11b+ F4/80+ | 0.02 | 0.9413 |
| Candidatus Saccharimonas | % CD45+ CD11b+ F4/80+ | 0.05 | 0.8677 |
| Flavonifractor | % CD45+ CD11b+ F4/80+ | -0.22 | 0.4876 |
| Parasutterella | % CD45+ CD11b+ F4/80+ | 0.09 | 0.7822 |
| Odoribacter | % CD45+ CD11b+ F4/80+ | -0.42 | 0.1731 |
| Mucispirillum | % CD45+ CD11b+ F4/80+ | -0.14 | 0.6598 |
| Oscillospira | % CD45+ CD11b+ F4/80+ | -0.33 | 0.2929 |
| Muribaculum | % CD45+ CD11b+ F4/80+ | 0.24 | 0.4572 |
| Herbinix | % CD45+ CD11b+ F4/80+ | 0.58 | 0.0503 |
| Negativibacillus | % CD45+ CD11b+ F4/80+ | 0.27 | 0.4034 |
| [Ruminococcus] gnavus group | % CD45+ CD11b+ F4/80+ | -0.12 | 0.7196 |
| Oscillibacter | % CD45+ CD11b+ F4/80+ | -0.28 | 0.3828 |
| Anaerotruncus | % CD45+ CD11b+ F4/80+ | -0.16 | 0.6133 |
| Alistipes | % CD45+ CD11b+ F4/80+ | -0.49 | 0.1036 |
| Prevotella_7 | % CD45+ CD11b+ F4/80+ | 0.14 | 0.6574 |
| Harryflintia | % CD45+ CD11b+ F4/80+ | -0.10 | 0.7629 |
| Anaeroplasma | % CD45+ CD11b+ F4/80+ | -0.12 | 0.7135 |
| Parabacteroides | % CD45+ CD11b+ F4/80+ | -0.25 | 0.4353 |
| UCG-001 | % CD45+ CD11b+ F4/80+ | 0.21 | 0.5094 |
| Butyricicoccus | % CD45+ CD11b+ F4/80+ | 0.29 | 0.3580 |
| Stomatobaculum | % CD45+ CD11b+ F4/80+ | -0.54 | 0.0723 |
| Prevotellaceae NK3B31 group | % CD45+ CD11b+ F4/80+ | 0.14 | 0.6606 |
| Lactobacillus | % CD45+ CD11b+ F4/80+ | 0.19 | 0.5522 |
| [Eubacterium] eligens group | % CD45+ CD11b+ F4/80+ | -0.11 | 0.7328 |
| Angelakisella | % CD45+ CD11b+ F4/80+ | 0.36 | 0.2469 |
| ASF356 | % CD45+ CD11b+ F4/80+ | -0.15 | 0.6388 |
| [Eubacterium] nodatum group | % CD45+ CD11b+ F4/80+ | 0.26 | 0.4060 |
| Roseburia | % CD45+ CD11b+ F4/80+ | -0.12 | 0.7059 |
| Erysipelatoclostridium | % CD45+ CD11b+ F4/80+ | 0.30 | 0.3366 |
| Paludicola | % CD45+ CD11b+ F4/80+ | 0.05 | 0.8848 |
| Rikenella | % CD45+ CD11b+ F4/80+ | -0.65 | 0.0216 |
| Lachnospiraceae UCG-001 | % CD45+ CD11b+ F4/80+ | 0.28 | 0.3766 |
| Tyzzerella | % CD45+ CD11b+ F4/80+ | -0.28 | 0.3833 |
| **Supplementary Table S5B:** Correlation between immune cells population in TAV and Genus with differential abundance among diets. Correlation significant established using the Pearson test (Significant p-value <0.05 was consider).  Abbreviations: CD: cluster of differentiation, VAT: visceral adipose tissue, MFI: median fluorescence intensity. | | | |

**Supplementary Table S5C:** Correlation between immune cells population in TAV and blood metabolites

| **Supplementary Table S5C** | | | |
| --- | --- | --- | --- |
| **Immune cells variable in TAV** | **Blood metabolites** | **rho** | **p-value** |
| % CD45+ CD4+ | Glucose | -0.04 | 0.9128 |
| % CD45+ CD4+ FOXP3+ | Glucose | 0.59 | 0.0415 |
| % CD45+ CD4+ FOXP3+ IL-10+ | Glucose | -0.79 | 0.0021 |
| MFI (CD45+ CD4+ FOXP3+ IL-10) | Glucose | -0.67 | 0.0164 |
| % CD45+ CD11b+ | Glucose | 0.46 | 0.1327 |
| % CD45+ CD11b+ F4/80+ | Glucose | -0.09 | 0.7781 |
| % CD45+ CD4+ | Cholesterol | 0.02 | 0.9390 |
| % CD45+ CD4+ FOXP3+ | Cholesterol | 0.80 | 0.0017 |
| % CD45+ CD4+ FOXP3+ IL-10+ | Cholesterol | -0.74 | 0.0058 |
| MFI (CD45+ CD4+ FOXP3+ IL-10) | Cholesterol | -0.51 | 0.0875 |
| % CD45+ CD11b+ | Cholesterol | 0.15 | 0.6516 |
| % CD45+ CD11b+ F4/80+ | Cholesterol | -0.47 | 0.1260 |
| % CD45+ CD4+ | Triglycerides | -0.22 | 0.4859 |
| % CD45+ CD4+ FOXP3+ | Triglycerides | 0.52 | 0.0824 |
| % CD45+ CD4+ FOXP3+ IL-10+ | Triglycerides | -0.12 | 0.7024 |
| MFI (CD45+ CD4+ FOXP3+ IL-10) | Triglycerides | 0.09 | 0.7792 |
| % CD45+ CD11b+ | Triglycerides | 0.09 | 0.7913 |
| % CD45+ CD11b+ F4/80+ | Triglycerides | -0.20 | 0.5276 |
| % CD45+ CD4+ | Total.proteins | -0.30 | 0.3414 |
| % CD45+ CD4+ FOXP3+ | Total.proteins | 0.16 | 0.6240 |
| % CD45+ CD4+ FOXP3+ IL-10+ | Total.proteins | -0.40 | 0.1931 |
| MFI (CD45+ CD4+ FOXP3+ IL-10) | Total.proteins | 0.06 | 0.8456 |
| % CD45+ CD11b+ | Total.proteins | -0.04 | 0.8971 |
| % CD45+ CD11b+ F4/80+ | Total.proteins | -0.68 | 0.0160 |
| % CD45+ CD4+ | Albumin | 0.05 | 0.8758 |
| % CD45+ CD4+ FOXP3+ | Albumin | -0.03 | 0.9241 |
| % CD45+ CD4+ FOXP3+ IL-10+ | Albumin | -0.31 | 0.3276 |
| MFI (CD45+ CD4+ FOXP3+ IL-10) | Albumin | 0.09 | 0.7812 |
| % CD45+ CD11b+ | Albumin | -0.03 | 0.9215 |
| % CD45+ CD11b+ F4/80+ | Albumin | -0.37 | 0.2432 |
| % CD45+ CD4+ | AST | 0.25 | 0.4242 |
| % CD45+ CD4+ FOXP3+ | AST | -0.16 | 0.6155 |
| % CD45+ CD4+ FOXP3+ IL-10+ | AST | 0.35 | 0.2648 |
| MFI (CD45+ CD4+ FOXP3+ IL-10) | AST | 0.56 | 0.0596 |
| % CD45+ CD11b+ | AST | 0.53 | 0.0744 |
| % CD45+ CD11b+ F4/80+ | AST | -0.20 | 0.5249 |
| % CD45+ CD4+ | ALT | -0.16 | 0.6233 |
| % CD45+ CD4+ FOXP3+ | ALT | 0.71 | 0.0098 |
| % CD45+ CD4+ FOXP3+ IL-10+ | ALT | -0.39 | 0.2053 |
| MFI (CD45+ CD4+ FOXP3+ IL-10) | ALT | -0.32 | 0.3167 |
| % CD45+ CD11b+ | ALT | 0.08 | 0.8028 |
| % CD45+ CD11b+ F4/80+ | ALT | -0.13 | 0.6958 |
| % CD45+ CD4+ | Leptin | 0.21 | 0.5387 |
| % CD45+ CD4+ FOXP3+ | Leptin | 0.75 | 0.0080 |
| % CD45+ CD4+ FOXP3+ IL-10+ | Leptin | -0.56 | 0.0756 |
| MFI (CD45+ CD4+ FOXP3+ IL-10) | Leptin | -0.61 | 0.0474 |
| % CD45+ CD11b+ | Leptin | 0.34 | 0.3117 |
| % CD45+ CD11b+ F4/80+ | Leptin | 0.00 | 0.9983 |
| % CD45+ CD4+ | Adiponectin | -0.30 | 0.3700 |
| % CD45+ CD4+ FOXP3+ | Adiponectin | -0.32 | 0.3304 |
| % CD45+ CD4+ FOXP3+ IL-10+ | Adiponectin | 0.41 | 0.2113 |
| MFI (CD45+ CD4+ FOXP3+ IL-10) | Adiponectin | 0.60 | 0.0494 |
| % CD45+ CD11b+ | Adiponectin | 0.07 | 0.8351 |
| % CD45+ CD11b+ F4/80+ | Adiponectin | -0.39 | 0.2398 |
| **Supplementary Table S5C:** Correlation between immune cells population in TAV and blood metabolites. Correlation significant established using the Pearson test (Significant p-value <0.05 was consider).  Abbreviations: CD: cluster of differentiation, VAT: visceral adipose tissue, AST: aspartate aminotransferase, ALT: alanine aminotransferase, MFI: median fluorescence intensity. | | | |

**Supplementary Table S5D:** Correlation between blood metabolites and genus with differential abundance among diets

| **Supplementary Table 5D** |  |  |  |
| --- | --- | --- | --- |
| **Genus** | **Blood metabolites** | **rho** | **p-value** |
| Alloprevotella | Glucose | -0.62 | 0.0322 |
| Prevotellaceae UCG-001 | Glucose | -0.55 | 0.0658 |
| Lachnoanaerobaculum | Glucose | 0.29 | 0.3619 |
| Helicobacter | Glucose | 0.73 | 0.0071 |
| Candidatus Saccharimonas | Glucose | -0.27 | 0.3936 |
| Flavonifractor | Glucose | 0.06 | 0.8545 |
| Parasutterella | Glucose | -0.45 | 0.1401 |
| Odoribacter | Glucose | 0.47 | 0.1211 |
| Mucispirillum | Glucose | 0.84 | 0.0006 |
| Oscillospira | Glucose | 0.32 | 0.3122 |
| Muribaculum | Glucose | -0.35 | 0.2601 |
| Herbinix | Glucose | -0.23 | 0.4799 |
| Negativibacillus | Glucose | 0.29 | 0.3637 |
| [Ruminococcus] gnavus group | Glucose | 0.53 | 0.0796 |
| Oscillibacter | Glucose | 0.32 | 0.3133 |
| Anaerotruncus | Glucose | 0.84 | 0.0007 |
| Alistipes | Glucose | 0.22 | 0.4980 |
| Prevotella_7 | Glucose | -0.86 | 0.0003 |
| Harryflintia | Glucose | 0.03 | 0.9152 |
| Anaeroplasma | Glucose | -0.72 | 0.0087 |
| Parabacteroides | Glucose | -0.60 | 0.0401 |
| UCG-001 | Glucose | -0.24 | 0.4520 |
| Butyricicoccus | Glucose | -0.39 | 0.2140 |
| Stomatobaculum | Glucose | 0.21 | 0.5177 |
| Prevotellaceae NK3B31 group | Glucose | -0.30 | 0.3513 |
| Lactobacillus | Glucose | -0.34 | 0.2847 |
| [Eubacterium] eligens group | Glucose | 0.82 | 0.0011 |
| Angelakisella | Glucose | 0.55 | 0.0640 |
| ASF356 | Glucose | -0.23 | 0.4813 |
| [Eubacterium] nodatum group | Glucose | -0.57 | 0.0513 |
| Roseburia | Glucose | 0.81 | 0.0014 |
| Erysipelatoclostridium | Glucose | -0.56 | 0.0596 |
| Paludicola | Glucose | -0.20 | 0.5321 |
| Rikenella | Glucose | 0.29 | 0.3644 |
| Lachnospiraceae UCG-001 | Glucose | -0.36 | 0.2538 |
| Tyzzerella | Glucose | 0.36 | 0.2450 |
| Alloprevotella | Cholesterol | -0.74 | 0.0058 |
| Prevotellaceae UCG-001 | Cholesterol | -0.72 | 0.0088 |
| Lachnoanaerobaculum | Cholesterol | 0.22 | 0.4899 |
| Helicobacter | Cholesterol | 0.27 | 0.3980 |
| Candidatus Saccharimonas | Cholesterol | -0.16 | 0.6186 |
| Flavonifractor | Cholesterol | -0.08 | 0.8078 |
| Parasutterella | Cholesterol | -0.48 | 0.1183 |
| Odoribacter | Cholesterol | 0.53 | 0.0764 |
| Mucispirillum | Cholesterol | 0.37 | 0.2430 |
| Oscillospira | Cholesterol | 0.72 | 0.0084 |
| Muribaculum | Cholesterol | -0.63 | 0.0269 |
| Herbinix | Cholesterol | -0.47 | 0.1245 |
| Negativibacillus | Cholesterol | 0.19 | 0.5480 |
| [Ruminococcus] gnavus group | Cholesterol | 0.70 | 0.0120 |
| Oscillibacter | Cholesterol | 0.51 | 0.0902 |
| Anaerotruncus | Cholesterol | 0.73 | 0.0071 |
| Alistipes | Cholesterol | 0.53 | 0.0793 |
| Prevotella_7 | Cholesterol | -0.72 | 0.0084 |
| Harryflintia | Cholesterol | 0.24 | 0.4498 |
| Anaeroplasma | Cholesterol | -0.66 | 0.0206 |
| Parabacteroides | Cholesterol | 0.00 | 0.9894 |
| UCG-001 | Cholesterol | -0.45 | 0.1463 |
| Butyricicoccus | Cholesterol | -0.54 | 0.0705 |
| Stomatobaculum | Cholesterol | 0.23 | 0.4629 |
| Prevotellaceae NK3B31 group | Cholesterol | -0.38 | 0.2268 |
| Lactobacillus | Cholesterol | -0.40 | 0.1922 |
| [Eubacterium] eligens group | Cholesterol | 0.58 | 0.0503 |
| Angelakisella | Cholesterol | 0.11 | 0.7422 |
| ASF356 | Cholesterol | 0.26 | 0.4092 |
| [Eubacterium] nodatum group | Cholesterol | -0.63 | 0.0286 |
| Roseburia | Cholesterol | 0.53 | 0.0763 |
| Erysipelatoclostridium | Cholesterol | -0.59 | 0.0415 |
| Paludicola | Cholesterol | -0.21 | 0.5136 |
| Rikenella | Cholesterol | 0.76 | 0.0042 |
| Lachnospiraceae UCG-001 | Cholesterol | -0.43 | 0.1654 |
| Tyzzerella | Cholesterol | 0.67 | 0.0172 |
| Alloprevotella | Triglycerides | -0.26 | 0.4190 |
| Prevotellaceae UCG-001 | Triglycerides | -0.33 | 0.2995 |
| Lachnoanaerobaculum | Triglycerides | 0.39 | 0.2129 |
| Helicobacter | Triglycerides | -0.53 | 0.0794 |
| Candidatus Saccharimonas | Triglycerides | 0.10 | 0.7686 |
| Flavonifractor | Triglycerides | 0.18 | 0.5711 |
| Parasutterella | Triglycerides | -0.19 | 0.5600 |
| Odoribacter | Triglycerides | 0.40 | 0.1962 |
| Mucispirillum | Triglycerides | -0.26 | 0.4182 |
| Oscillospira | Triglycerides | 0.15 | 0.6445 |
| Muribaculum | Triglycerides | -0.45 | 0.1416 |
| Herbinix | Triglycerides | 0.03 | 0.9200 |
| Negativibacillus | Triglycerides | 0.33 | 0.2896 |
| [Ruminococcus] gnavus group | Triglycerides | 0.63 | 0.0288 |
| Oscillibacter | Triglycerides | 0.31 | 0.3294 |
| Anaerotruncus | Triglycerides | -0.10 | 0.7454 |
| Alistipes | Triglycerides | 0.41 | 0.1809 |
| Prevotella_7 | Triglycerides | -0.13 | 0.6865 |
| Harryflintia | Triglycerides | 0.69 | 0.0124 |
| Anaeroplasma | Triglycerides | -0.13 | 0.6847 |
| Parabacteroides | Triglycerides | 0.38 | 0.2177 |
| UCG-001 | Triglycerides | -0.37 | 0.2354 |
| Butyricicoccus | Triglycerides | -0.08 | 0.8020 |
| Stomatobaculum | Triglycerides | 0.58 | 0.0494 |
| Prevotellaceae NK3B31 group | Triglycerides | -0.34 | 0.2743 |
| Lactobacillus | Triglycerides | -0.08 | 0.8138 |
| [Eubacterium] eligens group | Triglycerides | -0.21 | 0.5150 |
| Angelakisella | Triglycerides | 0.02 | 0.9499 |
| ASF356 | Triglycerides | 0.39 | 0.2099 |
| [Eubacterium] nodatum group | Triglycerides | -0.11 | 0.7434 |
| Roseburia | Triglycerides | -0.16 | 0.6224 |
| Erysipelatoclostridium | Triglycerides | -0.06 | 0.8501 |
| Paludicola | Triglycerides | 0.10 | 0.7463 |
| Rikenella | Triglycerides | 0.55 | 0.0634 |
| Lachnospiraceae UCG-001 | Triglycerides | -0.33 | 0.2973 |
| Tyzzerella | Triglycerides | 0.71 | 0.0101 |
| Alloprevotella | Total.proteins | -0.20 | 0.5382 |
| Prevotellaceae UCG-001 | Total.proteins | -0.66 | 0.0189 |
| Lachnoanaerobaculum | Total.proteins | -0.56 | 0.0605 |
| Helicobacter | Total.proteins | 0.13 | 0.6832 |
| Candidatus Saccharimonas | Total.proteins | 0.31 | 0.3228 |
| Flavonifractor | Total.proteins | 0.32 | 0.3149 |
| Parasutterella | Total.proteins | 0.16 | 0.6274 |
| Odoribacter | Total.proteins | 0.09 | 0.7735 |
| Mucispirillum | Total.proteins | 0.27 | 0.3984 |
| Oscillospira | Total.proteins | 0.46 | 0.1296 |
| Muribaculum | Total.proteins | 0.13 | 0.6872 |
| Herbinix | Total.proteins | -0.80 | 0.0017 |
| Negativibacillus | Total.proteins | -0.48 | 0.1140 |
| [Ruminococcus] gnavus group | Total.proteins | 0.06 | 0.8608 |
| Oscillibacter | Total.proteins | 0.23 | 0.4745 |
| Anaerotruncus | Total.proteins | 0.30 | 0.3414 |
| Alistipes | Total.proteins | 0.36 | 0.2477 |
| Prevotella_7 | Total.proteins | -0.33 | 0.2913 |
| Harryflintia | Total.proteins | 0.04 | 0.9107 |
| Anaeroplasma | Total.proteins | 0.12 | 0.7054 |
| Parabacteroides | Total.proteins | 0.20 | 0.5294 |
| UCG-001 | Total.proteins | 0.17 | 0.5960 |
| Butyricicoccus | Total.proteins | -0.05 | 0.8833 |
| Stomatobaculum | Total.proteins | 0.23 | 0.4768 |
| Prevotellaceae NK3B31 group | Total.proteins | -0.24 | 0.4433 |
| Lactobacillus | Total.proteins | 0.27 | 0.3958 |
| [Eubacterium] eligens group | Total.proteins | 0.15 | 0.6436 |
| Angelakisella | Total.proteins | -0.47 | 0.1258 |
| ASF356 | Total.proteins | 0.30 | 0.3491 |
| [Eubacterium] nodatum group | Total.proteins | -0.03 | 0.9290 |
| Roseburia | Total.proteins | 0.35 | 0.2619 |
| Erysipelatoclostridium | Total.proteins | -0.41 | 0.1842 |
| Paludicola | Total.proteins | 0.34 | 0.2729 |
| Rikenella | Total.proteins | 0.51 | 0.0911 |
| Lachnospiraceae UCG-001 | Total.proteins | -0.59 | 0.0437 |
| Tyzzerella | Total.proteins | 0.26 | 0.4202 |
| Alloprevotella | Albumin | 0.03 | 0.9322 |
| Prevotellaceae UCG-001 | Albumin | -0.30 | 0.3369 |
| Lachnoanaerobaculum | Albumin | -0.69 | 0.0121 |
| Helicobacter | Albumin | 0.04 | 0.9079 |
| Candidatus Saccharimonas | Albumin | 0.34 | 0.2855 |
| Flavonifractor | Albumin | 0.43 | 0.1611 |
| Parasutterella | Albumin | 0.17 | 0.6068 |
| Odoribacter | Albumin | -0.16 | 0.6223 |
| Mucispirillum | Albumin | 0.13 | 0.6880 |
| Oscillospira | Albumin | 0.66 | 0.0186 |
| Muribaculum | Albumin | 0.21 | 0.5161 |
| Herbinix | Albumin | -0.88 | 0.0002 |
| Negativibacillus | Albumin | -0.72 | 0.0089 |
| [Ruminococcus] gnavus group | Albumin | 0.08 | 0.7996 |
| Oscillibacter | Albumin | 0.48 | 0.1133 |
| Anaerotruncus | Albumin | 0.26 | 0.4069 |
| Alistipes | Albumin | 0.09 | 0.7812 |
| Prevotella_7 | Albumin | -0.04 | 0.9065 |
| Harryflintia | Albumin | 0.18 | 0.5732 |
| Anaeroplasma | Albumin | 0.15 | 0.6410 |
| Parabacteroides | Albumin | 0.44 | 0.1488 |
| UCG-001 | Albumin | 0.24 | 0.4569 |
| Butyricicoccus | Albumin | 0.21 | 0.5076 |
| Stomatobaculum | Albumin | 0.20 | 0.5332 |
| Prevotellaceae NK3B31 group | Albumin | -0.08 | 0.7958 |
| Lactobacillus | Albumin | 0.26 | 0.4052 |
| [Eubacterium] eligens group | Albumin | 0.12 | 0.7026 |
| Angelakisella | Albumin | -0.70 | 0.0115 |
| ASF356 | Albumin | 0.56 | 0.0596 |
| [Eubacterium] nodatum group | Albumin | 0.04 | 0.9140 |
| Roseburia | Albumin | 0.31 | 0.3261 |
| Erysipelatoclostridium | Albumin | -0.47 | 0.1215 |
| Paludicola | Albumin | 0.46 | 0.1303 |
| Rikenella | Albumin | 0.34 | 0.2822 |
| Lachnospiraceae UCG-001 | Albumin | -0.74 | 0.0064 |
| Tyzzerella | Albumin | 0.29 | 0.3562 |
| Alloprevotella | AST | -0.15 | 0.6516 |
| Prevotellaceae UCG-001 | AST | 0.32 | 0.3106 |
| Lachnoanaerobaculum | AST | -0.16 | 0.6120 |
| Helicobacter | AST | 0.37 | 0.2367 |
| Candidatus Saccharimonas | AST | -0.17 | 0.6026 |
| Flavonifractor | AST | -0.15 | 0.6373 |
| Parasutterella | AST | 0.04 | 0.8963 |
| Odoribacter | AST | -0.40 | 0.1961 |
| Mucispirillum | AST | 0.18 | 0.5847 |
| Oscillospira | AST | -0.22 | 0.4977 |
| Muribaculum | AST | 0.26 | 0.4228 |
| Herbinix | AST | 0.04 | 0.8934 |
| Negativibacillus | AST | -0.10 | 0.7564 |
| [Ruminococcus] gnavus group | AST | -0.38 | 0.2265 |
| Oscillibacter | AST | -0.17 | 0.5983 |
| Anaerotruncus | AST | 0.19 | 0.5474 |
| Alistipes | AST | -0.54 | 0.0723 |
| Prevotella_7 | AST | 0.29 | 0.3549 |
| Harryflintia | AST | 0.02 | 0.9591 |
| Anaeroplasma | AST | 0.16 | 0.6138 |
| Parabacteroides | AST | 0.21 | 0.5097 |
| UCG-001 | AST | 0.05 | 0.8869 |
| Butyricicoccus | AST | 0.25 | 0.4314 |
| Stomatobaculum | AST | 0.28 | 0.3854 |
| Prevotellaceae NK3B31 group | AST | 0.08 | 0.8058 |
| Lactobacillus | AST | -0.25 | 0.4327 |
| [Eubacterium] eligens group | AST | 0.24 | 0.4446 |
| Angelakisella | AST | 0.01 | 0.9765 |
| ASF356 | AST | 0.07 | 0.8394 |
| [Eubacterium] nodatum group | AST | 0.18 | 0.5801 |
| Roseburia | AST | 0.31 | 0.3336 |
| Erysipelatoclostridium | AST | 0.10 | 0.7643 |
| Paludicola | AST | -0.28 | 0.3709 |
| Rikenella | AST | -0.24 | 0.4430 |
| Lachnospiraceae UCG-001 | AST | 0.14 | 0.6670 |
| Tyzzerella | AST | -0.20 | 0.5290 |
| Alloprevotella | ALT | -0.49 | 0.1068 |
| Prevotellaceae UCG-001 | ALT | -0.61 | 0.0335 |
| Lachnoanaerobaculum | ALT | 0.26 | 0.4102 |
| Helicobacter | ALT | 0.12 | 0.7131 |
| Candidatus Saccharimonas | ALT | 0.28 | 0.3744 |
| Flavonifractor | ALT | 0.09 | 0.7860 |
| Parasutterella | ALT | -0.32 | 0.3074 |
| Odoribacter | ALT | 0.28 | 0.3866 |
| Mucispirillum | ALT | 0.37 | 0.2414 |
| Oscillospira | ALT | 0.44 | 0.1566 |
| Muribaculum | ALT | -0.48 | 0.1141 |
| Herbinix | ALT | -0.28 | 0.3766 |
| Negativibacillus | ALT | 0.41 | 0.1907 |
| [Ruminococcus] gnavus group | ALT | 0.58 | 0.0486 |
| Oscillibacter | ALT | 0.43 | 0.1655 |
| Anaerotruncus | ALT | 0.51 | 0.0876 |
| Alistipes | ALT | 0.34 | 0.2778 |
| Prevotella_7 | ALT | -0.63 | 0.0297 |
| Harryflintia | ALT | 0.54 | 0.0686 |
| Anaeroplasma | ALT | -0.60 | 0.0379 |
| Parabacteroides | ALT | 0.19 | 0.5478 |
| UCG-001 | ALT | -0.25 | 0.4288 |
| Butyricicoccus | ALT | -0.21 | 0.5155 |
| Stomatobaculum | ALT | 0.30 | 0.3457 |
| Prevotellaceae NK3B31 group | ALT | -0.41 | 0.1843 |
| Lactobacillus | ALT | -0.28 | 0.3818 |
| [Eubacterium] eligens group | ALT | 0.33 | 0.3001 |
| Angelakisella | ALT | 0.18 | 0.5802 |
| ASF356 | ALT | 0.57 | 0.0523 |
| [Eubacterium] nodatum group | ALT | -0.29 | 0.3656 |
| Roseburia | ALT | 0.49 | 0.1022 |
| Erysipelatoclostridium | ALT | -0.47 | 0.1252 |
| Paludicola | ALT | -0.08 | 0.7932 |
| Rikenella | ALT | 0.54 | 0.0686 |
| Lachnospiraceae UCG-001 | ALT | -0.40 | 0.1960 |
| Tyzzerella | ALT | 0.83 | 0.0008 |
| Alloprevotella | Leptin | -0.66 | 0.0286 |
| Prevotellaceae UCG-001 | Leptin | -0.38 | 0.2515 |
| Lachnoanaerobaculum | Leptin | 0.76 | 0.0069 |
| Helicobacter | Leptin | 0.45 | 0.1652 |
| Candidatus Saccharimonas | Leptin | -0.21 | 0.5277 |
| Flavonifractor | Leptin | -0.23 | 0.5020 |
| Parasutterella | Leptin | -0.59 | 0.0573 |
| Odoribacter | Leptin | 0.36 | 0.2698 |
| Mucispirillum | Leptin | 0.42 | 0.1948 |
| Oscillospira | Leptin | 0.07 | 0.8286 |
| Muribaculum | Leptin | -0.68 | 0.0226 |
| Herbinix | Leptin | 0.18 | 0.5921 |
| Negativibacillus | Leptin | 0.72 | 0.0132 |
| [Ruminococcus] gnavus group | Leptin | 0.56 | 0.0743 |
| Oscillibacter | Leptin | 0.05 | 0.8899 |
| Anaerotruncus | Leptin | 0.64 | 0.0339 |
| Alistipes | Leptin | 0.11 | 0.7467 |
| Prevotella_7 | Leptin | -0.72 | 0.0125 |
| Harryflintia | Leptin | 0.42 | 0.2017 |
| Anaeroplasma | Leptin | -0.85 | 0.0009 |
| Parabacteroides | Leptin | -0.47 | 0.1423 |
| UCG-001 | Leptin | -0.49 | 0.1227 |
| Butyricicoccus | Leptin | -0.73 | 0.0108 |
| Stomatobaculum | Leptin | 0.03 | 0.9209 |
| Prevotellaceae NK3B31 group | Leptin | 0.04 | 0.9134 |
| Lactobacillus | Leptin | -0.51 | 0.1062 |
| [Eubacterium] eligens group | Leptin | 0.61 | 0.0445 |
| Angelakisella | Leptin | 0.70 | 0.0166 |
| ASF356 | Leptin | -0.15 | 0.6655 |
| [Eubacterium] nodatum group | Leptin | -0.49 | 0.1225 |
| Roseburia | Leptin | 0.52 | 0.1013 |
| Erysipelatoclostridium | Leptin | 0.05 | 0.8946 |
| Paludicola | Leptin | -0.45 | 0.1650 |
| Rikenella | Leptin | 0.22 | 0.5213 |
| Lachnospiraceae UCG-001 | Leptin | -0.11 | 0.7515 |
| Tyzzerella | Leptin | 0.38 | 0.2477 |
| Alloprevotella | Adiponectin | -0.07 | 0.8316 |
| Prevotellaceae UCG-001 | Adiponectin | 0.01 | 0.9690 |
| Lachnoanaerobaculum | Adiponectin | -0.37 | 0.2685 |
| Helicobacter | Adiponectin | -0.18 | 0.5888 |
| Candidatus Saccharimonas | Adiponectin | -0.13 | 0.7067 |
| Flavonifractor | Adiponectin | 0.53 | 0.0960 |
| Parasutterella | Adiponectin | -0.02 | 0.9486 |
| Odoribacter | Adiponectin | 0.13 | 0.6979 |
| Mucispirillum | Adiponectin | 0.07 | 0.8303 |
| Oscillospira | Adiponectin | 0.03 | 0.9292 |
| Muribaculum | Adiponectin | 0.34 | 0.3138 |
| Herbinix | Adiponectin | -0.08 | 0.8241 |
| Negativibacillus | Adiponectin | -0.40 | 0.2186 |
| [Ruminococcus] gnavus group | Adiponectin | -0.19 | 0.5802 |
| Oscillibacter | Adiponectin | 0.34 | 0.3137 |
| Anaerotruncus | Adiponectin | -0.20 | 0.5608 |
| Alistipes | Adiponectin | 0.16 | 0.6342 |
| Prevotella_7 | Adiponectin | 0.40 | 0.2244 |
| Harryflintia | Adiponectin | -0.03 | 0.9237 |
| Anaeroplasma | Adiponectin | 0.53 | 0.0904 |
| Parabacteroides | Adiponectin | 0.53 | 0.0958 |
| UCG-001 | Adiponectin | 0.15 | 0.6571 |
| Butyricicoccus | Adiponectin | 0.48 | 0.1388 |
| Stomatobaculum | Adiponectin | 0.66 | 0.0270 |
| Prevotellaceae NK3B31 group | Adiponectin | -0.45 | 0.1624 |
| Lactobacillus | Adiponectin | 0.01 | 0.9801 |
| [Eubacterium] eligens group | Adiponectin | -0.09 | 0.7954 |
| Angelakisella | Adiponectin | -0.37 | 0.2619 |
| ASF356 | Adiponectin | 0.29 | 0.3901 |
| [Eubacterium] nodatum group | Adiponectin | 0.11 | 0.7520 |
| Roseburia | Adiponectin | -0.22 | 0.5219 |
| Erysipelatoclostridium | Adiponectin | -0.34 | 0.3107 |
| Paludicola | Adiponectin | 0.35 | 0.2882 |
| Rikenella | Adiponectin | -0.01 | 0.9769 |
| Lachnospiraceae UCG-001 | Adiponectin | -0.06 | 0.8529 |
| Tyzzerella | Adiponectin | 0.01 | 0.9865 |
| **Supplementary Table S5D:** Correlation between blood metabolites and genus with differential abundance among diets. Correlation significant established using the Pearson test (Significant p-value <0.05 was consider).  Abbreviations: AST: aspartate aminotransferase, ALT: alanine aminotransferase. | | | |

**Supplementary Table S6:** displays the values derived from the mediation model analysis. It outlines the results for distinct regression pathways (a, b, c) (see Figure 12), as well as the indirect and total effects. For each mediation involving the predicted variable (metabolic parameter), predictor variable (microbial genus), and mediator variable (immunological parameter), standardized estimates, upper and lower confidence intervals, and p-values are included.
